# MJF-14 proximity ligation assay detects early non-inclusion alpha-synuclein pathology with enhanced specificity and sensitivity

**DOI:** 10.1101/2024.07.08.602186

**Authors:** Nanna Møller Jensen, YuHong Fu, Cristine Betzer, Hongyun Li, Sara Elfarrash, Ali H. Shaib, Donatus Krah, Zagorka Vitic, Lasse Reimer, Hjalte Gram, Vladimir Buchman, Mark Denham, Silvio O. Rizzoli, Glenda M. Halliday, Poul Henning Jensen

## Abstract

Lewy pathology, consisting of Lewy bodies and Lewy neurites, is the pathological hallmark of synucleinopathies such as Parkinson’s disease and dementia with Lewy bodies, but it is generally thought to represent late-stage pathological changes. In contrast, α-synuclein oligomers are regarded as early-stage pathology, likely involved in disease progression and cellular toxicity. Oligomers, however, are not detected by standard immunohistochemistry but require specific detection techniques such as the proximity ligation assay (PLA). Here, we describe the MJF-14 PLA, a new PLA towards aggregated α-synuclein with unprecedented specificity, attained by the utilization of aggregate conformation-specific α-synuclein antibody MJFR-14-6-4-2 (hereafter MJF-14). Signal in the assay directly correlates with α-synuclein aggregation in SH-SY5Y cells, as treatment with aggregation inhibitor ASI1D significantly lowers PLA signal. In human cortical neurons, MJF-14 PLA detects pre-formed fibril-induced aggregation, especially prominent when using stealth PFFs invisible to the MJF-14 antibody. Co-labelling of MJF-14 PLA and pS129-α-synuclein immunofluorescence in post-mortem dementia with Lewy bodies cases showed that while the MJF-14 PLA reveals extensive non-inclusion pathology, it is not sensitive towards Lewy bodies. In Parkinson’s disease brain, direct comparison of PLA and IHC with the MJF-14 antibody, combined with machine learning-based quantification, showed striking α-synuclein pathology preceding the formation of conventional Lewy pathology. The majority of the PLA-revealed non-inclusion pathology was found in the neuropil, including some clearly located in the presynaptic terminals. With this work, we introduce an improved α-synuclein aggregate PLA to uncover abundant non-inclusion pathology, which deserves future validation with multiple brain bank resources and in different synucleinopathies.

## Introduction

Since the identification of aggregated α-synuclein as a primary constituent of the Lewy bodies (LBs) and Lewy neurites, histopathological hallmarks of certain neurological diseases, now known as synucleinopathies, researchers have sought to elucidate the exact involvement of α-synuclein aggregates in the pathogenesis of these diseases. Comprising Parkinson’s disease (PD), dementia with Lewy bodies (DLB), and multiple system atrophy (MSA) as the most prominent, the synucleinopathies are principally age-related neurodegenerative diseases ^1–3^. Although the majority of cases are sporadic, specific mutations and mul-tiplications of the SNCA gene cause familial PD, underscoring the importance of α-synuclein in the path-ogenesis ^4–9^. In certain conditions, α-synuclein readily misfolds and accumulates into various aggregates; first the small soluble oligomers and later the larger, insoluble fibrils, which can then associate with other cellular components and form LBs ^10–12^. Mapping of Lewy pathology in PD patients with variable disease duration has revealed distinct patterns of pathology in the central nervous system, leading to the prion-like spreading hypothesis ^13–16^ and one of the current classifications of disease severity: the Braak staging ^17^.

Though the Lewy bodies are striking markers of synucleinopathies, it has long been contested whether the LBs actually cause neuronal death and confer the spreading of pathology between cells – or whether they represent protective cellular responses sequestering aberrant α-synuclein, or are simply a by-stander phenomenon ^11,18–22^. As a consequence, attention has fallen upon the α-synuclein oligomers, which are much smaller, soluble structures formed during the aggregation process ^23–32^. Oligomers are thus hypothesized to be a major culprit in both disease symptomatology and progression of synucle-inopathies – as well as a potential therapeutic target ^33–39^. However, due to their smaller size, detecting oligomeric structures is challenging. In standard immunohistochemistry, the potential presence of oligo-mers is dwarfed by the presence of abundant native α-synuclein and the much larger, intensely stained LBs and Lewy neurites, and the specificity of truly oligomer-selective antibodies remains questionable, although antibodies with oligomer-preference have been reported ^40,41^.

As an alternative to the oligomer-specific antibodies, proximity-dependent strategies have been em-ployed to detect oligomers, including the bimolecular fluorescence complementation assay (BiFC) and the proximity ligation assay (PLA). Where the BiFC utilizes a proximity-dependent completion of a fluoro-phore to detect the early stages of α-synuclein aggregation ^42,43^, the PLA relies on antibody-based bind-ing of proximity probes within approximately 40 nm, followed by a signal amplification to achieve high sensitivity ^44^. In 2015, using the syn211 antibody against total α-synuclein, Roberts et al. demonstrated a preferential staining of oligomers over both fibrils and monomeric α-synuclein in the so far predominant α-synuclein PLA; in PD patient tissue, the PLA mainly stained early-stage pathology preceding the Lewy bodies, such as pale bodies ^45^. The use of a total α-synuclein-antibody, however, can pose an issue, es-pecially in settings where α-synuclein expression is increased, as the PLA in itself does not preclude the detection of closely located monomers of the protein, e.g. when bound to cellular vesicles.

In this paper, we describe the development of a new α-synuclein PLA with excellent specificity obtained by the utilization of conformation-specific antibody MJFR-14-6-4-2 (MJF-14) towards aggregated α-synuclein. We directly demonstrated this specificity in a cell model overexpressing α-synuclein, in conditions with and without α-synuclein aggregation, and compared it against the existing syn211 PLA. Furthermore, we characterized the usability of the MJF-14 PLA, both in fluorescent and chromogenic applications, in cultured human cortical neurons, post-mortem brain tissue of PD and DLB, and in mouse models of synucleinopathies. Lastly, we described the development of a machine-learning-based automated segmentation and quantification of chromogenic PLA, enabling large-scale quantification studies.

## Materials and methods

### Protein production, purification and fibril assembly

Full-length S129A-mutated human α-synuclein and S129A-mutated human α-synuclein with an added glycine residue at the C-terminal (S129A-α-synuclein-141G) was expressed in BL21(DE3)-competent cells and purified by reverse phase chromatography as previously described ^46–48^. To prepare pre-formed fibrils (PFFs), soluble monomeric α-synuclein (4 mg/mL) was incubated in PBS at 37°C with 1050 rpm shaking for 72 hours, before collection, characterization, and storage as previously described ^48–50^. Immediately before use, aliquots of PFFs were quickly thawed, sonicated briefly using a Branson SFX 250 sonifier (0.3 s on, 0.7 s off, 30% power, for 45 pulses). Sonicated aliquots were stored at room temperature until use.

### Cell culture

Human neuroblastoma (SH-SY5Y) cells with inducible expression of α-synuclein ^51^ were grown in Roswell Park Memorial Institute (RPMI) 1640 medium (Lonza, #BE12-702F) supplemented with 15% fetal bovine serum (Biowest, #S1810), 50 units/mL penicillin and 50 μg/mL streptomycin (Merck, #A2213), 200 µg/mL geneticin (G418, TCI EUROPE, #TCIAG0349), 50 µg/mL hygromycin B (Gibco, #10687010), and 1 µg/mL doxycycline (Calbiochem, #324385). On day 0, cells were seeded onto coverslips and α-synuclein expression was induced in some cultures by the removal of doxycycline. For all experiments, mitotic SH-SY5Y cells were differentiated into non-mitotic neuronal cells by the addition of 20 µM all-trans retinoic acid on day 1 (Molecular Probes/Invitrogen, #207340010). From day 1 onwards, 20 µM of ASI1D was added to inhibit aggregation of α-synuclein in some of the cultures (> 98% pure H-RGGAVVTGRRRRRR-NH_2_ (Schafer-N), 10 mM stock in 98% ethanol) ^52^. At 7 days post expression of α-synuclein, cells were fixed in 4% PFA.

### Human cortical neurons

Human embryonic stem cells (hESCs) NN3053 (Novo Nordisk) were differentiated to cortical neurons in a similar manner to what has been previously described ^53,54^. Briefly, NN3053 hESCs were plated on Lam-inin-521 in mTeSR1 medium with 10 µM Rock inhibitor Y-27632 (Tocris Bioscience). The following day, media was changed to CPN medium (1:1 DMEM/F12 and Neurobasal with 0.5x B27, 0.5x N2, 0.5x ITSA, 1x GlutaMAX, 0.5x penicillin/streptomycin, and 50 µM 2-mercaptoethanol (Life Technologies)) and supplemented with 100 nM LDN193189 (Stemgent) and 10 mM SB431542 (Tocris Bioscience). Media was changed every second day or as required. On day 11 neural progenitors were passaged with gentle dissociation buffer (PBS supplemented with 0.5 mM EDTA) and reseeded on laminin-521 coated plates at a split ratio of 1:2.5 in CPN medium supplemented with 20 ng/mL fibroblast growth factor 2 (FGF2; R&D Systems) and 10 µM Y-27632. The following day, media were changed to CPN basal media containing 20 ng/mL FGF2. On day 18 cells were passaged with gentle dissociation buffer and reseeded at 60,000 cells/cm^2^ on laminin-521 in CPN medium supplemented with 10 µM Y-27632. The following day media was changed to CPN medium and from then changed as required. On day 25, cells were plated on Laminin-521 coated plates at 90,000 cells/cm^2^ in CPN basal media supplemented with 10 µM Y-27632, 200 nM ascorbic acid (AA; Sigma-Aldrich), 0.05 mM dcAMP (Sigma-Aldrich), 40 ng/mL GDNF (R&D systems). The following day and onwards, media was changed to Pan-neuronal medium (1:1 DMEM/F12 and Neurobasal with 0.5x B27, 0.5x N2, 1x GlutaMAX, 0.5x penicillin/streptomycin, and 1x NEAA (Life Technologies)) supplemented with 200 nM AA, 0.05 mM dcAMP, 40 ng/mL GDNF, 2 µM LM22A4 (Tocris Bioscience), 2.5 µM DAPT (Tocris Bioscience), and 1 ug/mL laminin (Sigma-Aldrich).

On day 35, Pan-neuronal medium with supplements was prepared with 42 µg/mL of either S129A or S129A-α-synuclein-141G (stealth) PFFs, or a similar volume of sterile PBS. One third of the medium was replaced in each well, resulting in a final concentration of PFFs of 14 µg/mL. Neurons were fixed at either 2 hours or 7 days post treatment, with the regular media change schedule resumed after 24 hours of treatment. After fixation in 4% PFA, cultures were dried completely to increase adhesion to glass slides, before addition of PBS and storage at 4°C until staining. Cultures were subjected to either a combined PLA and immunofluorescence (IF) protocol or to IF only.

### Animal tissue

All procedures involving animals were conducted in accordance with the European directive on animal experiments (2010/63/EU), and the studies were approved by the Danish Animal Experiments Inspectorate (license no. 2017-15-0201-01203). All mice were housed in a temperature-controlled room under 12 h light/dark cycles with unlimited access to food and water.

To study the usability of MJF-14 PLA in mouse models, we employed a range of different mouse strains. These included wild-type C57Bl6 mice (WT), α-synuclein transgenic mice (line 61; B6;DBA-Tg(Thy1-SNCA)61Ema ^55^) overexpressing human α-synuclein (ASO, males only), two different strains of α-synuclein knockout mice (ASKO #1; SNCA^−/−^; C57BL/6N-Snca^tm1Mjff^/J; JAX stock #016123 ^56^, and ASKO #2; SNCA^−/−^; B6(Cg)-*Snca*^tm1.2Vlb^/J; JAX stock #028559 ^57,58^), α-/β-synuclein double knockout mice (A/B-KO) and α-/β-/*γ*-synuclein triple knockout mice (TKO). A/B-KO and TKO mice were generated as described by Connor-Robson et al., albeit using the ASKO strain #2 for α-synuclein knockout ^59^. All mice studied were adult (approx. 4-13 months old) and we studied 2-4 mice of each strain. The total number of experimental mice used in this study was 17.

Mice were sacrificed by intraperitoneal injection of lethal dosage of pentobarbital and perfused with PBS followed by 4% PFA, both containing phosphatase inhibitors (25 mM β-glycerolphosphate, 5 mM NaF, 1 mM Na_3_VO_4_, and 10 mM Na-pyrophosphate) before removing the brain. Brains were post-fixed in 4% PFA for 24 h and then stored in PBS with 0.05% sodium azide before paraffin-embedding and microtome sectioning at 8-10 µm.

### Human tissue

Brain samples from pathologically confirmed subjects with PD (n = 10 early PD; n = 10 late PD) and controls without any neurological or neuropathological disease (n = 10) were obtained from the Sydney Brain Bank (see Table 1). Controls were matched for age (average 77-80 y) and post-mortem delay (average 9-15 h). All cases with PD were levodopa-responsive and met the UK Brain Bank Clinical Criteria for a diagnosis of PD with no other neurodegenerative conditions. PD and control cases were assessed according to the regional presence of Lewy pathology (Braak *et al.*, 2003), with all controls having no Lewy pathology and the PD cases having Braak stages IV (early PD) or VI (late PD) Lewy pathology. The mean disease duration did not differ between the early (average 13 ± 2 y) and late PD (average 14 ± 8 y) groups (see Table 1). The study was approved by the University of Sydney Human Research Ethics Committee (2017/985).

**Table 1.** Demography of the human anterior cingulate cortex cohort (data presented as mean ± SD)

|  | Control | Early PD (Braak IV) | Late PD (Braak VI) | P-value |
| --- | --- | --- | --- | --- |
| Sex (M/F) | 5/5 | 5/5 | 6/4 | 0.88 ($\chi^2$) |
| Age (years) | 78±10 | 77±6 | 80±4 | 0.57 (ANOVA) |
| Post-mortem delay (hours) | 15±6 | 14±8 | 9±10 | 0.22 (ANOVA) |
| Disease duration (years) | - | 13±2 | 14±8 | 0.32 (T) |

As we have shown previously the anterior cingulate cortex (ACC) has staged α-synuclein changes in PD with increased levels but limited Lewy inclusions at stage IV progressing to significant Lewy inclusions by stage VI ^60^, we requested formalin-fixed paraffin-embedded (FFPE) tissue sections from the ACC for this study.

To assess the ability of the PLA to detect Lewy bodies and assess neuropil PLA localization, FFPE sections from the superior frontal cortex and motor cortex of patients with dementia with Lewy bodies (DLB) (n = 2) and from non-neurodegenerative controls (n = 2) were similarly obtained from the Sydney Brain Bank. Average age was 82 years (range 80-84 y) and post-mortem delay in the range 9.5-14.5 h. Both DLB cases were Braak Lewy stage 6, with Braak tangle stage 0-1 and mild-moderate CERAD neuritic plaques, while the controls had no Lewy or significant age-related pathologies.

FFPE sections were cut on a rotary microtome (Thermo/Microm, #HM325) at 8 µm and mounted on Series 2 adhesive slides (Trajan Scientific Medical).

### Antibody conjugation for proximity ligation assay

Proximity ligation assay (PLA) was carried out using either the Duolink PLA kits for brightfield (Sigma, #DUO92012) or fluorescent applications (Sigma, #DUO92008), NaveniBright HRP (Navinci, #NB.MR.HRP.100) for brightfield applications or a custom-made detection kit utilizing biotin in the fluorophore labelling process (similarly obtained from Navinci). Assays were performed using the confor-mation-specific α-synuclein antibody MJF-14 (MJFR-14-6-4-2; Abcam, #ab214033) or the total α-synuclein antibody syn211 (Abcam, #ab206675).

For Duolink PLA applications, antibodies were conjugated to plus-and minus-oligonucleotides using the Duolink PLA Probemaker kits (Sigma, #DUO92009 and #DUO92010) according to manufacturer’s protocol. In brief, 20 µL antibody was mixed with 2 µL conjugation buffer, added to the lyophilized oligonucleotides (plus-or minus-) and incubated overnight at room temperature (RT). 2 µL stop reagent was then added and incubated for 30 minutes at RT. Finally, 24 µL of storage solution was added and conjugated antibodies stored at 4°C.

For Navinci PLA applications, MJF-14 was conjugated to Navenibody 1 and 2 oligonucleotides using the NaveniLink conjugation kit (Navinci, #NL.050) according to manufacturer’s protocol. In brief, 10 µL of Modifier was added to 100 µL antibody, before mixing with the lyophilized oligonucleotides (Navenibody 1 or Navenibody 2) and incubated overnight at room temperature (RT). 10 µL of Quencher N was then added and incubated for 15 minutes at RT, and conjugated antibodies were stored at 4°C.

In general, Duolink PLA probes and kits were used in this manuscript, except where specifically stated that Navinci PLA probes and kits were used instead.

### Proximity ligation assay

The following general protocol was used to perform PLA, with slight variations in incubation times and antibody dilutions depending on cells/tissues and whether fluorescent or chromogenic PLA was performed. Exact incubation times and antibody dilutions can be found in Table 2. Unless otherwise mentioned, washing was performed in 1x TBS + 0.05% Tween-20. For paraffin sections, initial deparaffinization and rehydration steps were followed by heat-induced epitope retrieval in 1x citrate buffer at pH 6.1 (DAKO, #S1699). All samples were then permeabilized in 0.5% Triton X-100 in PBS, followed by endogenous peroxide quenching in 0.3% hydrogen peroxide (chromogenic PLA only). Samples were then blocked in 10% bovine serum albumin (BSA, Duolink MJF-14 PLA), 1x Duolink Blocking Solution (syn211 PLA), or 1x Naveni Blocking Buffer with Supplement 1 (Navinci MJF-14 PLA). For Duolink protocols, conjugated primary antibodies were diluted in 5% BSA (MJF-14 PLA) or in 1x Duolink Probe Diluent (syn211 PLA) and incubated for 1 hour at 37°C, followed by overnight incubation at 4°C. For Navinci MJF-14 PLA, conjugated primary antibodies were diluted in 1x Primary Antibody Diluent with Supplement 2 and incubated overnight at 4°C. Unbound antibody was washed off, and samples were incubated with Duolink ligation solution or Navinci Reaction 1 solution at 37°C, followed by washing and subsequent incubation in Duolink amplification solution or Navinci Reaction 2 solution at 37°C.

**Table 2.** Parameters for proximity ligation assay in various models. Unless otherwise specified, Duolink kits were used for the application.

| Cell or tissue | Staining method | Permeabilization, peroxide quenching and blocking times | Primary antibody dilution | Ligation time | Amplification time |
| --- | --- | --- | --- | --- | --- |
| **SH-SY5Y cells** | Fluorescent MJF-14 PLA | 30 minutes | 1:10,000 | 1 hour | 3 hours |
|  | Fluorescent syn211 PLA | 30 minutes (permeabilization)<br>1 hour (blocking) | 1:100 | 1 hour | 3 hours |
| **Human cortical neurons** | Fluorescent MJF-14 PLA | 30 minutes | 1:10,000 | 1 hour | 3 hours |
| **Human DLB and control FFPE** | Fluorescent MJF-14 PLA | 1 hour | 1:50,000 | 1.5 hours | 3.5 hours |
| **Human PD and control ACC FFPE** | Chromogenic MJF-14 PLA | 1 hour | 1:1000 | 1.5 hours | 3 hours |
| **Human DLB and control FFPE (expansion PLA)** | Fluorescent MJF-14 PLA (Navinci) | 30 minutes (permeabilization)<br>1 hour (blocking) | 1:5000 | 30 minutes | 60 minutes |
| **Mouse FFPE** | Chromogenic MJF-14 PLA (Duolink) | 1 hour | 1:10,000 / 1:25,000 | 30 minutes | 100 minutes |
|  | Chromogenic MJF-14 PLA (Navinci) | 1 hour | 1:10,000 | 30 minutes | 60 minutes |

For fluorescent Duolink PLA, the amplification process was followed by washing, another round of blocking in 10% BSA and regular immunostaining with either rat α-tubulin (Abcam, #ab6160, 1:1000) or mouse pS129 α-synuclein (11A5, kindly provided by Imago Pharmaceuticals, 1:25,000) and chicken neuronal βIII-tubulin (TUJ1, LS Bio, #LS-B5225, 1:500) in 5% BSA, overnight at 4°C. After washing off unbound antibody, samples were incubated for 3 hours at RT in secondary antibody (anti-rat AlexaFluor-488, #A21208, or anti-mouse AlexaFluor-488, Invitrogen #A11001, and anti-chicken AlexaFluor-647, Abcam #ab150175, as matching the primary antibodies) diluted 1:1000-1:2000 along with DAPI (4’,6-diamidino-2-phenylindole, TH.GEYER, 5 µg/mL) in 5% BSA, again followed by washing. Before mounting, autofluorescence in human post-mortem tissue was quenched by 30 seconds incubation in 1x TrueBlack in 70% ethanol (Biotium, #23007), followed by three washes in PBS. Then, samples were mounted with DAKO fluorescent mounting medium (DAKO, #S3023) and edges sealed with nail polish.

For chromogenic Duolink PLA, the amplification process was followed by washing and incubation in Duo-link detection solution for 1 hour at RT. After another round of washing, samples were incubated with Duolink substrate solution for 20 minutes at RT, washed again and nuclei counterstained with Duolink nuclear stain for 2 minutes at RT. Samples were left under running tap water for 10 minutes and then dehydrated in increasing concentrations of ethanol, incubated in NeoClear (Sigma, #109843), and mounted with DPX mounting medium (Sigma, #06522).

For chromogenic Navinci PLA, Reaction 2 was followed by washing twice in 1x TBS, then once in 0.1x TBS, and incubation in Navinci HRP reaction for 30 minutes at RT. After another round of washing in 1x TBS, samples were incubated with Navinci substrate reaction for 5 minutes, washed in dH_2_O and nuclei counterstained with Navinci nuclear stain for 5 seconds. Samples were left under running tap water for 10 minutes and then quickly dehydrated in 100% isopropanol and mounted with VectaMount Express Mounting Medium (Vector Laboratories, #H-5700).

In all experiments, technical negative controls (either PLA without primary conjugated antibodies or PLA without ligase in the ligation reaction) were included to confirm the specificity of the observed signal.

For experiments on transgenic mice, we additionally employed biological negative controls in the form of α-synuclein knockout mice (ASKO, 2 strains), α-/β-synuclein double knockout mice, and α-/β-/*γ*-synuclein triple knockout mice.

### Immunofluorescence (IF)

For IF of cultured human cortical neurons, all steps were performed in the chambered slides. Cultures were permeabilized in 0.5% Triton X-100, before blocking in 10% BSA and incubation with primary antibodies against aggregated α-synuclein (MJF-14; rabbit MJFR-14-6-4-2, Abcam, #ab209538, 1:25,000), total α-synuclein (mouse LB509, Abcam, #ab27766, 1:500), and neuronal βIII-tubulin (chicken TUJ1, LS Bio, #LS-B5225, 1:500) in 5% BSA for 1 hour at RT. Unbound antibody was washed off in 1x TBS + 0.05% Tween-20, followed by incubation with appropriate secondary antibodies (anti-rabbit AlexaFluor-488, Invitrogen #A11008, anti-mouse AlexaFluor-568, Invitrogen #A11004, and anti-chicken AlexaFluor-647, Abcam #ab150175) for 1 hour at RT. After another round of washing, most of the buffer was removed, plastic wells broken off the slides, and silicone gasket was removed with a pair of forceps to allow mounting with DAKO fluorescent mounting medium (DAKO, #S3023). After leaving slides to dry, edges were sealed with nail polish.

### Immunohistochemistry (IHC)

Before staining, all sections were deparaffinized and rehydrated with series of xylene and gradient ethanol. For IHC on human brain, the antigen was unmasked by 70% (vol/vol) formic acid (Sigma, #F0507-1L), followed by heat induced antigen retrieval in a programmable antigen retrieval cooker (Aptum Bio Retriever 2100) using 1X R-UNIVERSAL Epitope Recovery Buffer (EMS, #62719-10). The sections were then blocked with BLOXALL® Endogenous Alkaline Phosphatase Blocking Solution (VectorLabs, #SP-6000) followed by 5% horse serum before incubation with the primary antibody MJF-14 (anti-conformation-specific α-synuclein filament antibody MJFR-14-6-4-2, Abcam, #ab209538, 1:200). The immunohistology signals were visualized with ImmPRESS®-AP Horse Anti-Rabbit IgG Polymer (VectorLabs, #MP-5401) and Im-mPACT™ Vector®Red Alkaline Phosphatase Substrate (VectorLabs, #SK-5105) as per manufacturer’s instruction. After counterstaining with cresyl violet, sections were coverslipped with DPX mounting medium (Sigma, #06522).

For IHC on mouse brain, staining was performed in parallel with MJF-14 PLA with identical steps for epitope retrieval, permeabilization, and blocking of endogenous peroxidases. Sections were blocked in 5% normal goat serum (NGS) in PBS before incubation with the primary antibody MJF-14 (anti-confor-mation-specific α-synuclein filament antibody MJFR-14-6-4-2, Abcam, #ab209538, 1:25,000), overnight at 4°C. Biotinylated goat anti-rabbit IgG (VectorLabs, #BA-1000-1.5, 1:200) in 2.5% NGS was applied for 1 hour at RT, followed by Vectastain® Elite® ABC-HRP kit (VectorLabs, #PK-6100) for 30 minutes at RT. Staining was visualized with 3,3ʹ-diaminobenzidine (DAB; Sigma, #D5637) for 10 minutes at RT, and sections were counterstained with nuclear stain from Navinci PLA kit (Navinci, #NB.MR.HRP.100). Samples were left under running tap water for 10 minutes before rapid dehydration in 100% isopropanol and mounting with VectaMount Express Mounting Medium (Vector Laboratories, #H-5700).

### ONE microscopy of PLA

To apply ONE microscopy imaging ^61^ to MJF-14 PLA, the PLA protocol was modified slightly to accommodate the specific needs related to staining intensity for the expansion protocol and run with the custom-made PLA detection kit from Navinci, allowing fluorescent labelling of the PLA product using biotin. FFPE sections mounted on glass slides were treated as described above regarding deparaffinization, rehydration, epitope retrieval, permeabilization and PLA blocking steps. Then, a 2-step biotin blocking following manufacturer’s instructions (Ready Probes Biotin Blocking Solution (Invitrogen, #R37628) was included, before incubation with conjugated primary antibodies overnight. Naveni Reaction 1 (ligation) and Reaction 2 (amplification) were performed as indicated in Table 2. Excess reaction reagents were washed off in 1x TBS, followed by 0.1x TBS, as per manufacturer’s instructions, then washed in PBS, before autofluorescence quenching in 1x TrueBlack (Biotium, #23007) as described above.

Samples were blocked again, this time in 2.5% BSA + 2.5% NGS + 2.5% normal donkey serum (NDS) in PBS. This was followed by incubation with primary antibodies for immunofluorescence: guineapig VGLUT1 (Synaptic Systems, #135304, 1:200) and Star635P-conjugated α-synuclein nanobody ^62^ (NbSyn2-St635P, Nanotag custom-made, 10 µM, 1:150) for 1.5 hours at RT. Unbound antibody was washed off with 1% BSA + 1% NGS + 1% NDS in PBS, before incubation with appropriate secondary antibodies (streptavidin-AlexaFluor-488 (Invitrogen, #S32354, 1:200) and donkey anti-guineapig Cy3 (Dianova, #706-165-148, 1:200) for 1.5 hours at RT. After another three washes in 1% BSA + 1% NGS + 1% NDS in PBS, samples were washed twice in PBS. A third of each sample was then mounted in Mowiol as a non-expanded staining control, while the rest was processed for tissue expansion.

Expansion protocol was performed as described in Saal et al. ^63^ but with 0.3 mg/mL Acryloyl-X (Invitrogen, #A20770) in MES buffer (N-(morpholino)ethane sulfonic acid-based saline solution, 19.52 mg/mL MES + 87.66 mg/mL NaCl, pH 6). The lower pH enhances the penetration depth of the anchor ^64^. Samples were incubated with Acryloyl-X in MES overnight at 4°C, protected from light. Gelation solution, containing N,N-dimethylacrylamide (DMAA), sodium acrylate, potassium persulfate, and N,N,N’,N’-Tet-ramethylethylendiamine (TEMED), was mixed and purged with N_2_ as previously described ^63,65^. A gelation chamber, inspired by Truckenbrodt et al. ^65^, was created in a 15-cm TC dish, covered with tinfoil and placed upside down. Two stacks of two square coverslips each were affixed to a piece of parafilm, with 20 mm space in-between, using a little bit of nail polish. Once slightly dried, the parafilm/coverslips were flipped upside-down to stick it inside the lid of the TC dish, pressing down the parafilm and letting it dry completely. Finally, gelation chamber was covered with wet tissues along the edges. Gelation solution (approx. 200 µL/slide) was then added in the parafilm-covered space between the coverslips, and slides with tissue sections were placed upside-down on top, edges resting on the coverslips. If necessary, extra gelation solution was pipetted in from the side to ensure coverage of the tissue, before the gelation chamber was closed, and gels left to polymerize overnight at 23°C (14-18 hours).

Next day, gels were washed 5x 30 minutes in disruption buffer (100 mM Tris, 5% Triton X-100, 1% SDS, pH 8) and then autoclaved in fresh disruption buffer in a cold-start autoclave on a 30-minute program at 110-121°C. After allowing the gels to cool down slowly in the autoclave, the samples, which normally detach from their coverslips during either disruption buffer washing or autoclaving, were still attached to the glass slides. Thus, tissue was scraped off the glass slides using a razor blade before proceeding to the expansion step. Then, gels were transferred to 22 × 22 cm culture dishes and approx. 400 mL ddH_2_O added to each dish. Water was exchanged twice immediately and then every 30-45 minutes until gels were fully expanded. Gels were left overnight in ddH_2_O before imaging.

### Imaging and quantification

#### Cells and human neurons

Samples were imaged on a Zeiss AxioObserver 7 inverted fluorescent microscope with the Zen Blue 2.3 software and 20-30 images were taken per coverslip/well at X63 magnification, using DAPI and tubulin staining as a guidance for the presence of cells. PLA particles were quantified within tubulin-labelled cells, then normalized to the number of cells using Fiji (Fiji Is Just ImageJ, version 2.3.0/1.53t, National Institutes of Health). A minimum of 260 cells were analysed per condition per replicate.

#### Human DLB cohort

Samples were imaged at X63 magnification on a Zeiss AxioObserver 7 inverted fluorescent microscope with the Zen Blue 2.3 software. For total PLA analysis in DLB and control cases, 20 images were taken randomly per sample, exported as TIFF, background subtracted, PLA particles counted, and total PLA area computed in Fiji. PLA particle area was averaged per image.

To evaluate PLA signal in LB-positive and LB-negative neurons, large neurons were manually defined from the DAPI-staining, using nucleus size and morphology as guidance (see Suppl. Fig. 3 for examples). From merged images, each neuron was then outlined and the original grey-scale PLA-channel TIFF was duplicated for each neuron. Neurons were categorised as either LB-positive (a LB in close proximity to a large neuronal nucleus) or LB-negative (no LB in close proximity). LBs without obvious association to a neuronal nucleus were not included in analysis. PLA images were then filtered using Gaussian blur (sigma = 2) and thresholded using the RenyiEntropy method (threshold = 350). Signals above threshold and larger than 10 pixels^2^ (corresponding to a minimum diameter of approx. 250 nm) were counted as PLA and total area of PLA for each neuron was computed. For each image, the PLA area/neuron was averaged between LB-positive and LB-negative neurons to allow a pairwise comparison of PLA area in LB-positive neurons compared to neighbouring (i.e., in the same image) LB-negative neurons.

#### Human ACC cohort

All PLA-and IHC-labelled sections were auto-scanned at X40 on an Olympus VS120 Slide Scanner at the same settings. Orientation was adjusted using the VS-DESKTOP software (Olympus Soft Imaging Solutions GmbH, ver. 2.9 13753) to allow digital images to be cropped at the comparable anatomical locations with similarly sized regions of interest (OlyVIA 3.3; length × width: 510 μm × 495 μm, providing an area of 252,450 μm^2^). To assess whether PLA and IHC revealed similar or distinct pathological information, the laminar distribution of PLA positive signals and IHC-revealed α-synuclein deposit particles were quantified for both superficial grey matter (SGM, containing cortical layers I-III) and deep grey matter (DGM, containing cortical layers V-VI). The cropped OlyVIA VSI files were converted into full resolution TIFF files with Fiji for quantification to be completed by researchers blinded to the group sources of cases.

For the quantification of PLA-positive signal, we initially developed a classifier to automatically segment the images into their components (PLA signal, nuclei, and extranuclear area) using the Trainable Weka Segmentation plugin in Fiji ^66^, based on an optimization cohort consisting of 6 images. To minimize quantification variability from artificial effects (e.g., colour development), images were categorised into intensely and weakly stained depending on the counterstain intensity of haematoxylin (Suppl. Fig. 2A-B), allowing a separate classifier suitable for nuclei segmentation in each group. Classifier performance was evaluated by the measures sensitivity, precision, and accuracy (see Suppl. Fig. 2) and optimized for sensitivity, i.e. as high detection of the neuronal nuclei as possible. PLA signal segmentation was identical for intensely and weakly stained groups.

For subsequent analyses, images were automatically segmented into PLA signals, nuclei, and extranuclear area, after which PLA signals and nuclei were defined from the probability maps. To distinguish neuronal nuclei from those of glia, a minimum size of 350 pixel (36.7 µm^2^, corresponding to an approx. minimum diameter of 7 µm) was set for neuronal nuclei. Total PLA particles were then computed in the entire image, in the neuronal cell bodies (i.e. inside neuronal nuclei and within 3.5 µm radius of nuclei), and in the neuropil (including any signal in glial nuclei) and normalized to either the corresponding tissue area or the neuronal number.

For individual neuronal PLA analyses, neurons were analysed one at a time, and the PLA particles were computed inside the nucleus and in the assumed cytoplasmic region (within a 3.5 µm distance from the nucleus). Then, the neuron was assigned to one of nine semi-quantitative groups. The entire analysis workflow is outlined in Suppl. Fig. 3.

For IHC, the images were quantified for total α-synuclein deposit particles (particles with a minimum length of 2 µm and intensity >50 above background) and LBs (large inclusions with diameter > 5 μm with compact signal). For the deposit particles, images were colour split with “colour deconvolution” using vectors “H&E 2” ^67,68^. MJF-14 signal at colour 2 was then counted using the Cell Counter plugin in Fiji with brightness and contrast slightly adjusted. Any oval or round particle with the longest axis >2 µm and an intensity >50 above the background was defined as a deposit particle in this analysis. LBs were counted manually and confirmed independently by two researchers.

#### Expansion PLA

Confocal imaging of non-expanded staining control samples was done on an Abberior Expert line setup (Abberior Instruments) with an IX83 microscope (Olympus) and a 100 × oil immersion objective (UPLSAPO, 1.4 NA; Olympus). Excitation lines of 485 nm, 561 nm, and 640 nm were used for AlexaFluor-488, Cy3, and Star635P, respectively. ONE microscopy imaging was carried out on TCS SP5 STED microscope (Leica Microsystems, Wetzlar, Germany) as described previously ^61^. Briefly, using a resonant scanner at a speed of 8 kHz, 1500 or 2000 frames of images were taken in uni-directional *xyct* line scans using 633, 561, and 488 nm excitation lines filtered through acousto-optical tunable filters and detected HyD detectors. Each *xyct* image stack was acquired at an 8-bit depth with a pixel size of slightly less of 98 nm, and a format of 128 x 128 pixels. Image acquisitions were imported into the ONE plugin in Fiji to obtain the super-resolved images.

#### Mouse cohort

Entire X40 slide scans of coronal sections were obtained on an Olympus VS120 Slide Scanner with the VS-ASW imaging software. Image files in the VSI format were opened in QuPath v.0.3.2 ^69^ and regions for comparison were selected. Somatosensory cortex and hippocampus ROIs were selected around bregma -2.055 mm (range -1.755 to -2.355 mm), while midbrain ROIs were selected in the vicinity of the red nucleus, approx. bregma -3.455 mm (range -3.18 to -3.68 mm), based on the Allen mouse brain atlas ^70^. No analysis was performed.

### Statistics

All data were checked for normality using the Shapiro-Wilks test, and appropriate tests were selected based on normality as well as the number of groups to compare. In cell experiments, groups were compared using a Kruskal-Wallis one-way ANOVA followed by the Dunn post hoc test to account for multiple comparisons, as not all groups were normally distributed. For human cortical neurons, groups were compared using a two-way ANOVA followed by Tukey’s multiple comparison test, with treatment as one grouping variable and technical replicate no. as the other grouping variable. For comparison of PLA particles in DLB and control brain sections, groups were compared using a Mann-Whitney U-test (total PLA/image) or a Wilcoxon matched-pairs signed rank test (PLA in LB-positive and LB-negative neurons) as data did not pass the test for normality. All statistics were two-tailed, performed in GraphPad Prism 9, with the exception of the human ACC data.

For the human ACC, all statistical analyses were performed using the SPSS software (IBM, ver. 26). Differences between all groups in demographic variables (Table 1) were assessed using one-way ANOVA (age and post-mortem delay), Pearson chi-square (gender), and Student’s T-test for disease duration between PD groups. Two-factor univariate analyses (group by region) were used to assess whether there were any regional differences in the PLA signals quantified in the ACC of the different groups, covarying for age, post-mortem delay, and gender, followed by post hoc Bonferroni tests. A Spearman’s rho correlation was performed to identify significant associations between PLA density in neuronal and neuropil compartments, between the densities of PLA signals and IHC-revealed deposit particles, and between deposit particles and LBs specifically. For correlation between deposit particles and LBs, cases with no LBs in an ROI were excluded. To determine if the distribution of PLA signals varied within the neurons of the different groups, univariate analyses of the PLA levels in the neurons (nucleus, cytoplasm, and both) as well as outside the neuronal somas (in the neuropil) were performed, covarying for age, post-mortem delay, and gender, and again post hoc Bonferroni tests were used to identify any significant group differences.

All data are displayed as mean ± standard error of the mean (SEM), unless otherwise stated, and p-values are indicated as * p<0.05, ** p<0.01, *** p<0.001, **** p<0.0001. Graphs were generated in GraphPad Prism 9, while figures were created using Adobe Illustrator and BioRender (Publication license agreement UF26YL0C58).

## Results

### Development of an α-synuclein aggregation-specific proximity ligation assay

Proximity ligation is based on the antibody-facilitated detection of two targets in close proximity, whereby signal amplification allows visualization of each individual proximity interaction (Fig. 1A). Thus, the choice of antibodies becomes crucial in determining which targets are detected. As α-synuclein overexpression models are regularly used in the research of PD and related synucleinopathies, we surmised that proximity ligation assays utilizing antibodies binding total α-synuclein might be suboptimal due to potential detection of closely located native monomeric α-synuclein species in the overexpression systems, e.g. associated to membrane surfaces. We hypothesized that using an antibody that only recognize aggregated α-synuclein would perform superior to a PLA detecting all α-synuclein molecules in close proximity. To test the hypothesis, we used Tet-off SNCA transgenic human neuroblastoma cells (SH-SY5Y AS), in which α-synuclein overexpression in non-mitotic differentiated cells is induced by the removal of doxycycline, leading to the development of α-synuclein oligomers upon induction of α-synuclein expression ^51,71^. In addition to setting up conditions with and without α-synuclein expression (± doxycycline), in some cultures, α-synuclein aggregation inhibitor ASI1D ^52^ was added to prevent aggregation of the expressed α-synuclein (Fig. 1B). Then, PLA was conducted using either the total α-synuclein antibody syn211, as has previously been published ^45,72–74^ or the aggregate-specific α-synuclein antibody MJFR-14-6-4-2 (MJF-14), which should only detect pathologically misfolded α-synuclein.

**Figure 1:**
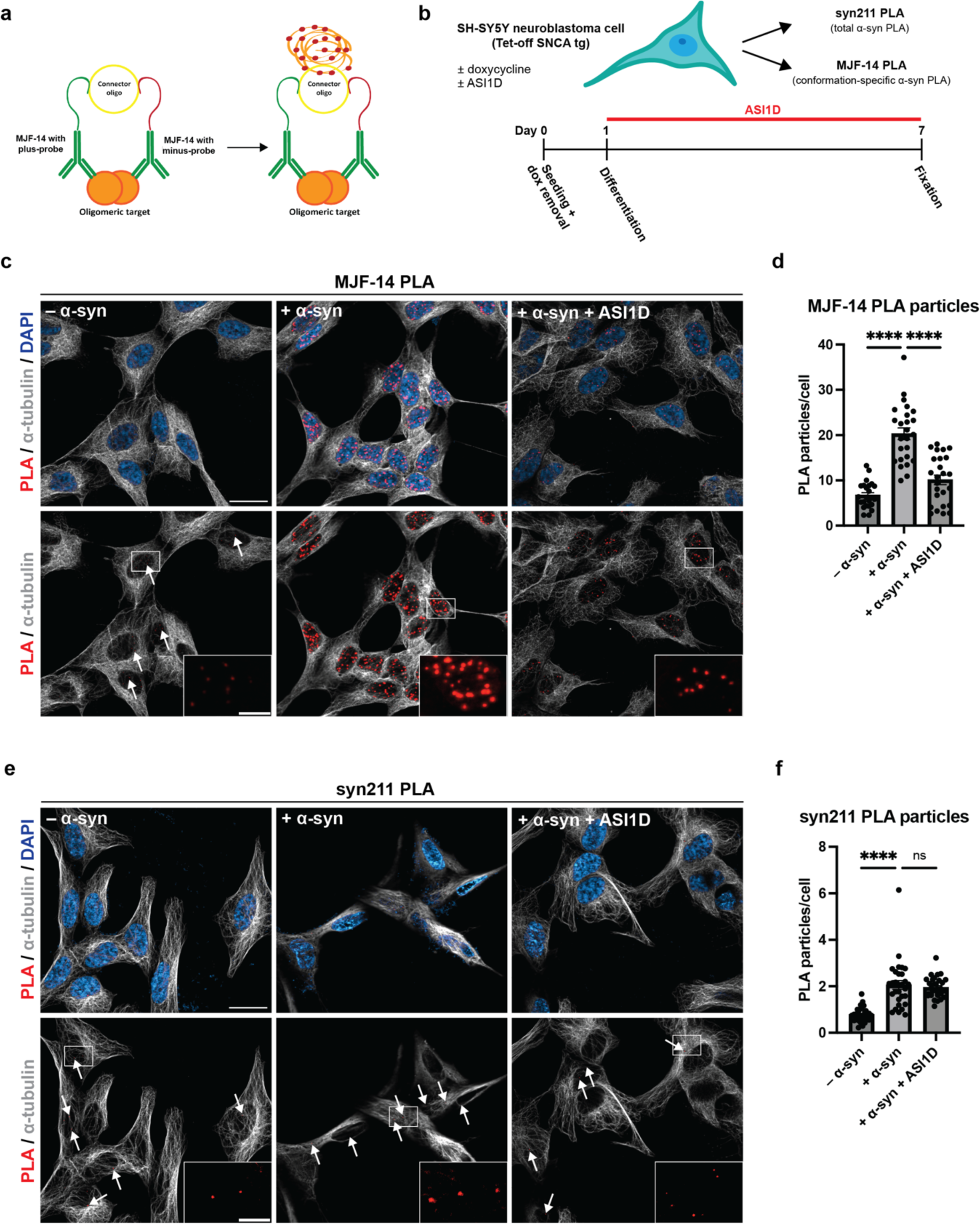
MJF-14 PLA signal is directly dependent upon aggregate formation in α-synuclein transgenic neuroblas-toma cells. **A.** Principle of the proximity ligation assay for detection of oligomeric targets. **B.** Experimental setup: α- synuclein transgenic SH-SY5Y cells are differentiated into non-mitotic neuronal-like cells using retinoic acid, treated ± doxycycline to mediate α-synuclein expression and ± ASI1D to modulate α-synuclein aggregation. Then, cells are fixed and subjected to PLA using the α-synuclein conformation-specific MJFR-14-6-4-2 antibody or the total α- synuclein syn211 antibody. **C.** Representative images of MJF-14 PLA in conditions without α-synuclein expression, with α-synuclein expression, and with α-synuclein expression + ASI1D-treatment. PLA signal is displayed in red (small dots), α-tubulin in grey and DAPI in blue. Arrows indicate examples of PLA particles in low abundance condi-tions. Scale bar = 20 µm. **D.** Quantification of the number of PLA particles per cell, as determined by the MJF-14 PLA. The +α-syn condition is significantly different from both -α-syn and +α-syn+ASI1D (p < 0.0001). **E.** Representa-tive images of syn211 PLA in conditions without α-synuclein expression, with α-synuclein expression, and with α- synuclein expression + ASI1D-treatment. PLA signal is displayed in red (small dots), α-tubulin in grey and DAPI in blue. Arrows indicate examples of PLA particles in low abundance conditions. Scale bar = 20 µm. **F.** Quantification of the number of PLA particles per cell, as determined by the syn211 PLA. The +α-syn condition is significantly dif-ferent from -α-syn (p < 0.0001) but not +α-syn+ASI1D (p > 0.9999). Graphs display mean ± SEM from one replicate and each dot signifies one image. Experiments were performed minimum three times independently, and groups were compared using a Kruskal-Wallis one-way ANOVA followed by the Dunn post hoc test. **** p<0.0001.

In conditions without α-synuclein overexpression, only a small amount of MJF-14 PLA signal was de-tected, while α-synuclein-overexpressing cultures contained widespread signal (Fig. 1C, D). Technical negative controls (PLA without ligase in the reaction) were included in all experiments, and their lack of signal for both antibodies ensured the functionality of the assay (data not shown). When the α-synuclein aggregate inhibitor ASI1D was added to the α-synuclein overexpressing cells, the MJF-14 PLA signal was returned almost to the same level as without α-synuclein overexpression (Fig. 1C, D). Of note, there were not only significantly more PLA particles in α-synuclein-overexpressing cultures than when treated with ASI1D or without α-synuclein overexpression, the particles were also significantly brighter (Fig. 1C, insets). A likely cause for this phenomenon is the presence of multiple rolling circle amplification prod-ucts in such a close vicinity that they coalesce in imaging, i.e., probably located on the same aggregate. Consequently, an even greater distinction between conditions could thus be achieved by quantifying the integrated density (i.e., intensity times area) of the PLA particles.

The syn211 PLA performed on parallel cultures also displayed an increase in PLA signal when α-synuclein overexpression was turned on but it was not reduced by ASI1D treatment (Fig. 1E, F). In general, the sig-nal from the syn211 PLA was much lower than that from the MJF-14 PLA, despite an increased antibody concentration (1:100 compared to 1:750 as previously published for fluorescent syn211 PLA) and efforts to optimize amplification time. This indicates that the MJF-14 PLA could be more sensitive towards very early types of aggregates. Collectively, we found that aggregation of α-synuclein is necessary for the generation of MJF-14 PLA signal, whereas the syn211 PLA may recognize physiological clustering as demonstrated, e.g., on surfaces of vesicles ^75–78^. Hence, we decided to further characterise the PLA based on the use of the MJF-14 antibody.

### MJF-14 PLA detects PFF-seeded pathology in human cortical neurons

In order to test the MJF-14 PLA in a different, likely more physiologically relevant, model, we cultured human cortical neurons in chambered slides and treated them with pre-formed fibrils (PFFs) to induce α-synuclein aggregation. Initial experiments showed a potential detection of the added PFFs by the MJF-14 PLA, especially at early timepoints, and we therefore included the novel stealth PFFs in our setup, invisible to the MJF-14 antibody due to a C-terminally added glycine, as recently described ^48^. Thus, cultured neurons were treated with either standard S129A-mutated PFFs or stealth PFFs (S129A-mutated α-synuclein-141G) and fixed at 2 hours or 7 days post treatment (Fig. 2A). At 2 hours post treatment, any signal above background levels in the cultures should stem from direct detection of the exogenous fibrils and, indeed, S129A PFF-treated cultures showed increased PLA signal (p < 0.0001, Fig. 2B-C). Signal in stealth PFF cultures, though, was not significantly different from the background level seen in PBS-treated cultures (p = 0.2318), indicating that the stealth PFFs were indeed not detected in the PLA. The same observation was made from MJF-14 IF of the cultures, whereas staining for total α-synuclein using the LB509 antibody showed similar, increased levels of α-synuclein after either type of PFF-treatment (Suppl. Fig. 1A). A closer examination of the IF images revealed signal deposition (both LB509 and MJF-14) not only along neuronal βIII-tubulin-positive processes but also in-between the neurons, signifying extracellular PFFs, of which only the S129A PFFs, but not the stealth PFFs, were detected by MJF-14 (Suppl. Fig. 1A). Staining for pS129-α-synuclein did not show any substantial signal at 2 hours post treatment, which was to be expected as both types of PFFs were S129A-mutated (Fig. 2B).

**Figure 2:**
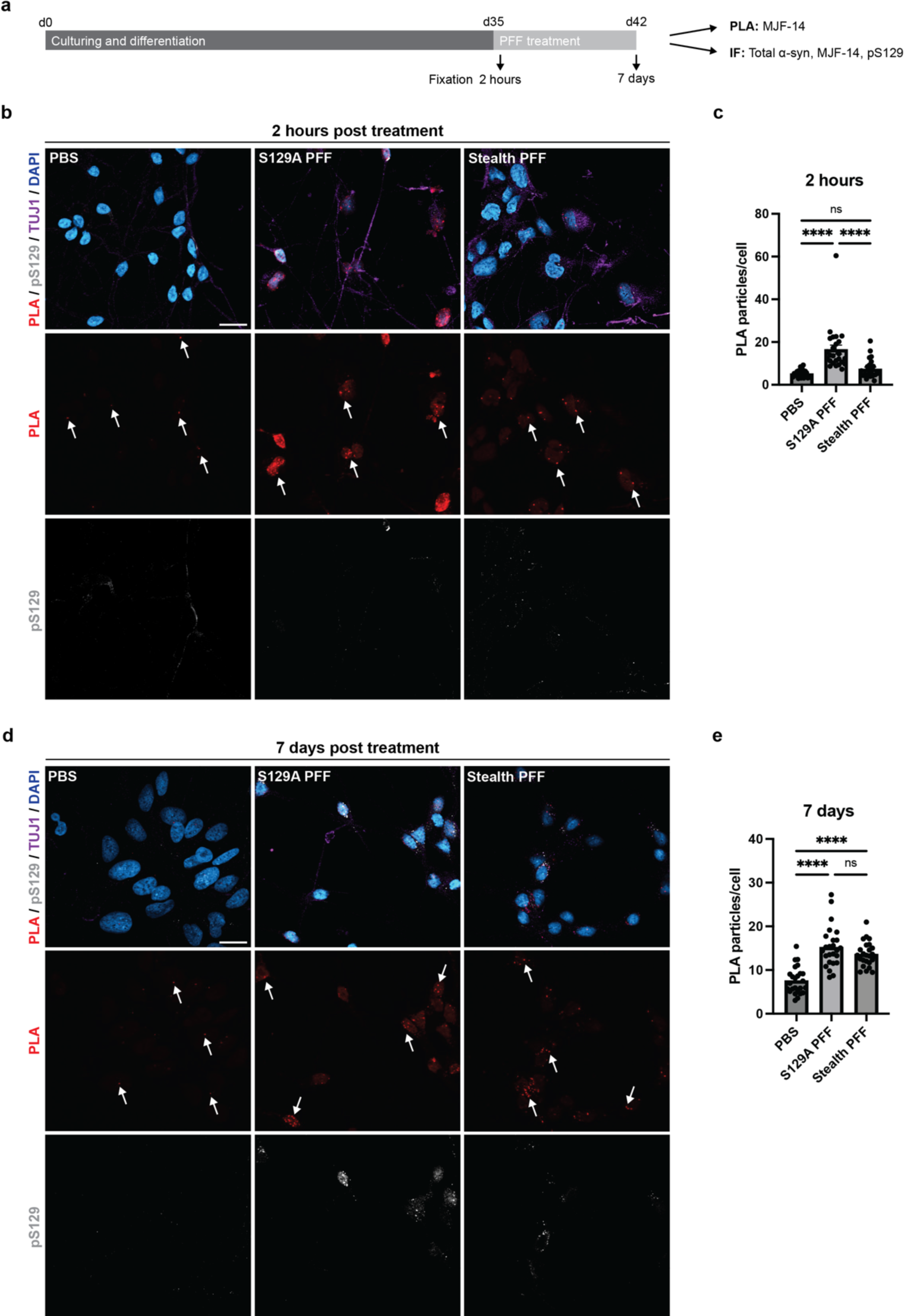
Stealth PFFs induce formation of PLA signal in human neurons but added PFFs themselves are unde-tected. **A.** Human cortical neurons were cultured for 35 days before treatment with either S129A PFFs, stealth PFFs (S129A-α-synuclein-141G), or PBS, and fixed after 2 hours or 7 days. **B.** Representative images from 2 hours post treatment cultures immunostained with MJF-14 PLA (red), pS129-α-synuclein (grey), βIII-tubulin/TUJ1 (purple), and DAPI nuclear stain (blue). Arrows indicate examples of PLA signals. Scale bar = 20 µm. **C.** Quantification of the number of PLA particles per cell after 2 hours, as determined by the MJF-14 PLA. Treatment significantly affects PLA signal density (F(2,269) = [47.79], p < 0.0001), with the S129A PFF group differing from both PBS and stealth PFF groups (p < 0.0001) while PBS and stealth PFF groups do no differ significantly (p = 0.2318). **D.** Representative images from 7 days post treatment cultures immunostained with MJF-14 PLA (red), pS129-α-synuclein (grey), βIII-tubulin (purple), and DAPI nuclear stain (blue). Arrows indicate examples of PLA signals. Scale bar = 20 µm**. E.** Quantification of the number of PLA particles per cell after 7 days, as determined by the MJF-14 PLA. Treatment significantly affects PLA signal density (F(2,296) = [35.26], p < 0.0001), with the PBS group differing from both S129A and stealth PFF groups (p < 0.0001) while S129A and stealth PFF groups do no differ significantly (p = 0.5009). Graphs display mean ± SEM from two independent replicates and each dot signifies one image. Experi-ments were performed two times independently with two technical replicates each time, and groups were com-pared using a two-way ANOVA followed by Tukey’s multiple comparison test. **** p<0.0001.

7 days after PFF treatment, internalized PFFs should have templated the aggregation of endogenous α- synuclein in the neurons, while the PFFs should be mostly truncated C-terminally and therefore undetectable by the MJF-14 antibody ^79–82^. As expected, treatment with both types of PFFs gave rise to increased MJF-14 PLA signal after 7 days (Fig. 2D-E), evincing aggregation of endogenous α-synuclein. This was further supported by IF detection of increased α-synuclein deposition following either PFF treatment, based on both pS129-α-synuclein (Fig. 2D) and MJF-14 IF (Suppl. Fig. 1B). Interestingly, although there was some co-detection of α-synuclein species by both the MJF-14 PLA and the pS129-α-synuclein IF, the two staining techniques also labelled distinct species, highlighting a discrepancy between phosphorylation and aggregation (Fig. 2D). We also observed that while α-synuclein accumulation into brightly fluorescent deposits was detected by MJF-14 after both PFF treatments, the signal-to-noise ratio was much higher in stealth PFF cultures (Suppl. Fig. 1B). This could indicate that a fraction of the S129A PFFs is still present, perhaps bound to the extracellular surface of the neurons, and thereby cause background staining.

Taken together, we show that the MJF-14 PLA can detect PFF-induced α-synuclein pathology but that it also has some detection of exogenous full-length PFFs. This challenge can be largely avoided by the use of stealth PFFs, which are invisible to the MJF-14 antibody and yield increased signal-to-noise ratio for both the MJF-14 PLA and regular MJF-14 IF.

### MJF-14 PLA reveals extensive non-Lewy body pathology in DLB brain samples

To further explore the MJF-14 PLA and its ability to stain synucleinopathy-related pathology, we studied sections of the superior frontal cortex from patients with dementia with Lewy bodies (DLB) and from brains without identified neurodegenerative disease. The sections were stained with the MJF-14 PLA in its fluorescent mode (red label) followed by a traditional immunofluorescence staining for pS129-α- synuclein (grey) to detect conventional Lewy body (LB) pathology. Initial staining of both DLB and con-trol brain sections revealed substantial fluorescence in the red channel, regardless of performing the PLA reaction with or without the antibodies added, and with or without the ligase in the PLA reaction (Fig. 3A). Most likely, the cause was lipofuscin-related autofluorescence, as described for human brain samples ^83^. Thus, an autofluorescence quenching step using TrueBlack was introduced, which efficiently suppressed autofluorescence and left the technical negative controls (PLA minus antibody and PLA mi-nus ligase) void of signal (Fig. 3B).

**Figure 3:**
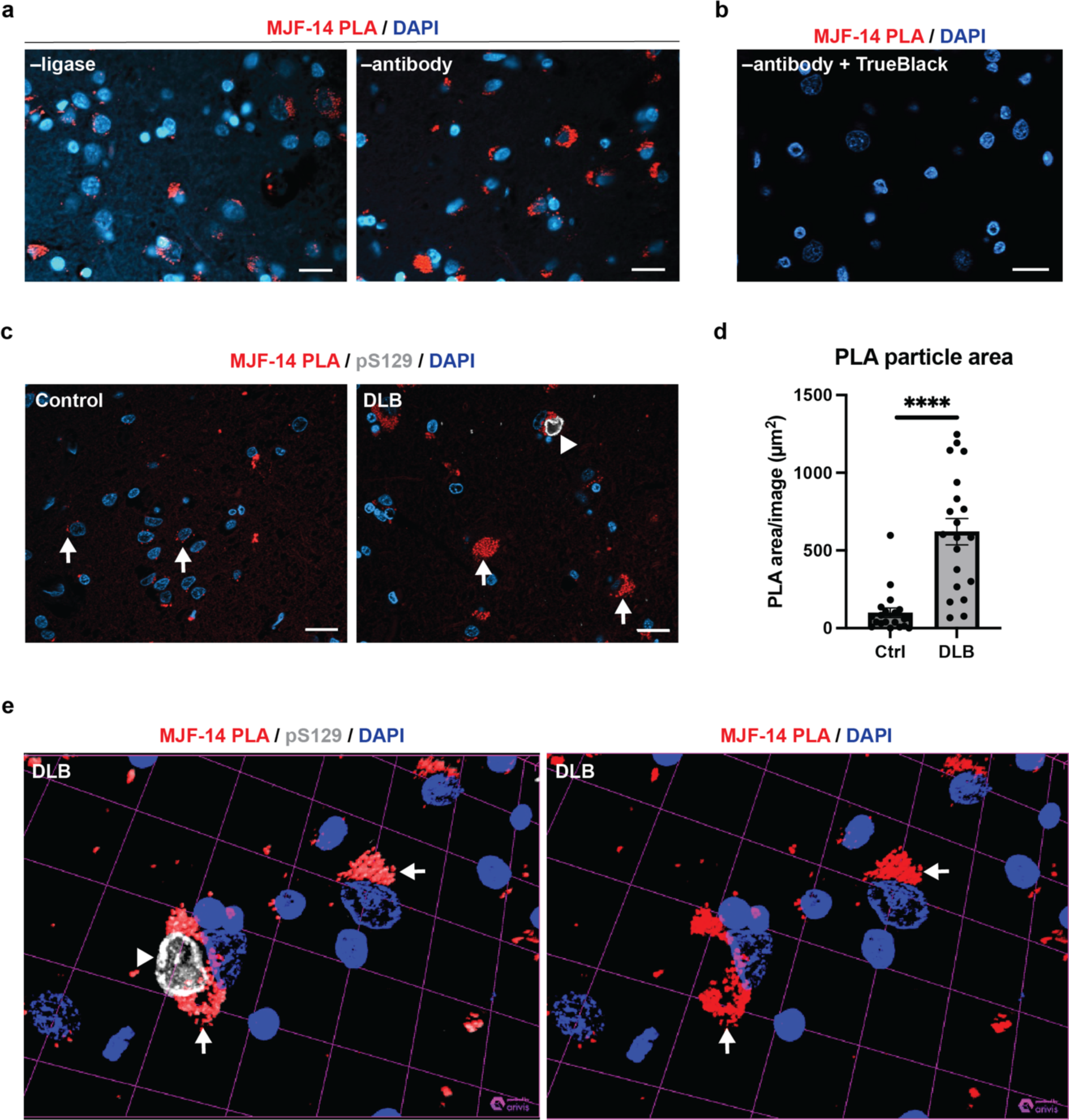
MJF-14 PLA stains considerable pathology in DLB brains but is not sensitive to Lewy bodies. **A.** Technical negative controls: PLA –ligase in the reaction (left) and PLA –antibody (right) display significant amounts of red channel signal when autofluorescence is not quenched. Scale bars = 20 µm. **B.** TrueBlack-quenching of autofluores-cence effectively removes background signal in technical negative controls (here, PLA –antibody). Scale bar = 20 µm. **C.** Representative images of sections immunostained with PLA (red), serine-129 phosphorylated α-synuclein (pS129, grey), and DAPI nuclear stain (blue) in control (left) and DLB (right). Examples of PLA-positive neurons are indicated with arrows, while a LB-positive neuron is indicated by an arrowhead. Scale bars = 20 µm. **D.** Quantifica-tion of PLA particle area in control and DLB, compared by a non-parametric Mann-Whitney U-test. Graph displays mean ± SEM of the total PLA area in µm^2^/image, with each dot representing one image. **** p<0.0001. **E.** Close-up z-stack rendering of a LB-containing neuron, with (left) and without (right) pS129, displaying the lack of PLA-stain-ing of the LB. PLA-signal (red, arrows) and LB (grey, arrowhead) is highlighted.

Following autofluorescence quenching, MJF-14 PLA-staining revealed considerable pathology in DLB brains, while control brains contained some PLA-staining, but at significantly lower levels as demonstrated by lower area covered by PLA per image (Fig. 3C, D). The level of PLA signal in non-neurodegenerative control brain was similar to what has previously been reported for the syn211 PLA on non-neurodegenerative controls by Roberts et al. ^45^. In the DLB brains, PLA-staining was found both in neurons with and without LBs, but there was only very sparse, if any, PLA-staining of the pS129-labelled LBs in the cases analysed (Fig. 3E). As the MJF-14 antibody does recognize Lewy bodies in IHC (see Fig. 4A), the inability of the MJF-14 PLA to efficiently label Lewy bodies is not antibody-related. Rather, the lack of PLA signal might indicate a structural organization of α-synuclein aggregates in LBs that differs from the MJF-14 PLA-positive oligomers and does not allow the necessary signal amplification in the proximity ligation assay.

**Figure 4:**
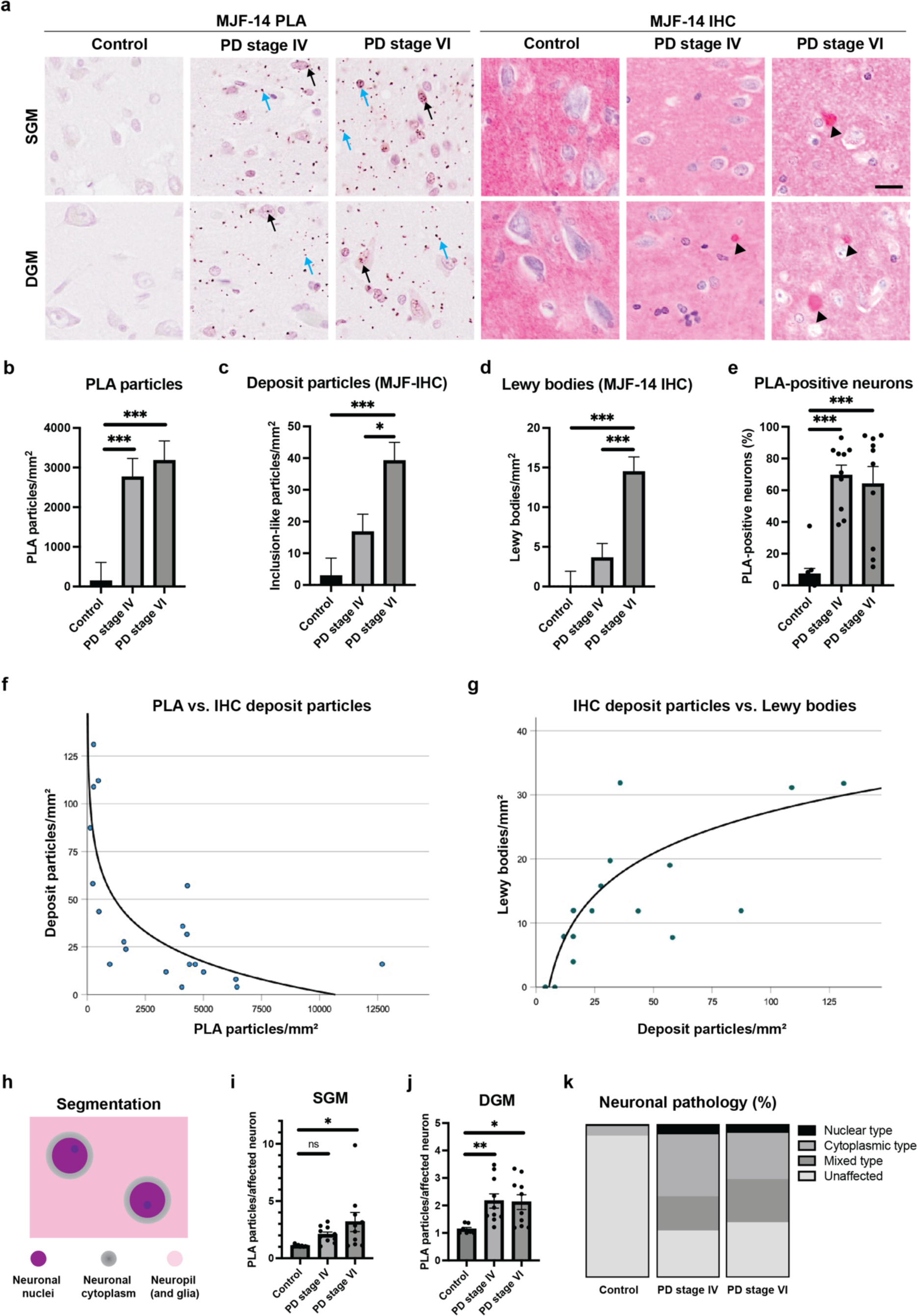
MJF-14 PLA efficiently distinguishes PD patients from controls but does not discriminate between early and late PD in the anterior cingulate cortex. **A.** Representative images comparing MJF-14 PLA and MJF-14 IHC in the superficial (SGM) and deep (DGM) layers of the ACC from control, early PD (Braak IV), and late PD (Braak VI). Black arrows indicate neuronal PLA, blue arrows indicate glial and neuropil PLA signals, and arrowheads highlight deposit particles/LBs. Scale bar = 20 µm. **B.** Quantification of the total PLA particle count per mm^2^. The control group is significantly different from the two PD groups (p<0.001). **C.** Quantification of the total deposit particle count per mm^2^ (detected by IHC). PD stage VI is significantly different from both controls (p<0.001) and PD stage IV (p=0.011). **D.** Quantification of LB count per mm^2^. PD stage VI is significantly different from both controls and PD IV (p<0.001). Values plotted in **B**-**D** can be found in Suppl. Table 2. **E.** Percentage of neurons containing PLA particles (averaged for SGM and DGM). Each subject is indicated as a dot. Both PD groups are significantly different from controls (p<0.001), but not from each other (p=0.409). For more information, see Suppl. Table 3. **F.** Correlation be-tween PLA particles and deposit particles (IHC) in PD stage VI. The density of PLA particles is inversely logarithmi-cally correlated with the density of deposit particles (r_s_ [20] = -0.732, p < 0.001), which fits a logarithmical model. **G**. Correlation between deposit particles and LBs in PD stage VI. The density of LBs is positively correlated with the density of deposit particles (r_s_ [20] = 0.675, p = 0.001). **H.** Schematic representation of image segmentation for analysis: PLA pathology is analysed inside neuronal cell bodies (in nuclei (purple) and/or assumed cytoplasm (grey)) as well as away from neuronal cell bodies (denoted neuropil, but also containing glia (pink)). **I.** Quantification of the average PLA particle count per affected neuron in SGM, with each subject indicated as one dot. There are sig-nificantly more PLA particles in affected neurons of PD stage VI than controls (p=0.011) and a similar non-signifi-cant tendency for PD stage IV (p=0.218). **J.** Quantification of the average PLA particle count per affected neuron in DGM, with each subject indicated as one dot. The control group is significantly different from both PD stage IV (p=0.007) and stage VI (p=0.011). Values plotted in **I**-**J** can be found in Suppl. Table 4. **K.** Proportion of different types of neuronal pathology, gradient grey coded for unaffected, solely nuclear, solely cytoplasmic, or mixed nu-clear and cytoplasmic types. Values used for generating the graphic can be found in Suppl. Table 5. Data are dis-played as mean ± SEM, averaged for SGM and DGM unless otherwise indicated, and comparisons are made using univariate analyses covarying with age, sex and post-mortem delay. * p<0.05, ** p<0.01, *** p<0.001.

### PD brains display significant MJF-14 PLA signal even at early stages

As chromogenic staining methods are often preferred by pathologists due to their stability, we tested the MJF-14 PLA in the chromogenic application on a cohort of Braak stage IV and VI PD patients and non-neurodegenerative controls (Table 1). For this, we selected the anterior cingulate cortex as our target region, as stage IV and VI PD patients are easily distinguishable by their Lewy pathology in this region (LBs are present in stage VI PD, but only rarely in stage IV PD) ^60^. A comparison of the MJF-14 PLA with regular MJF-14 IHC, i.e., using MJFR-14-6-4-2 antibody, on neighbouring sections clearly demonstrated the difference in specificity between conventional IHC and PLA. Where IHC preferentially detected Lewy bodies, the PLA uncovered abundant non-inclusion pathology distributed throughout the tissue of PD patients (Fig. 4A).

A disadvantage of chromogenic staining is the difficulty it poses in attempts to properly quantify pathology, as signals are not simply separated in different wavelength channels as for immunofluorescence. Instead, the image consists of many compound signals, which need to be separated before quantification. To do this, we employed a machine-learning based technique, the open-source Trainable Weka Segmentation in Fiji ^66^, to train a classifier, perform automated segmentation, and subsequently analyse the PLA-density in the sections (Suppl. Fig. 3).

Initially, we found striking PLA signal in PD, with no difference between stage IV and VI PD (p = 0.551). Both groups, however, were easily distinguished from the controls, which only contained very low levels of PLA signal (p < 0.001; Fig. 4B). As a comparison, we also quantified the MJF-14 IHC signal in the three groups, in the form of i) deposit particles (any MJF-14 signal with a minimum length of 2 µm and intensity >50 above background) and ii) LBs specifically (as identified by two independent researchers). As expected from the group selection criteria, MJF-14 IHC was not able to discriminate stage IV PD from controls by any of the two measures (p = 0.093 and p = 0.169, respectively; Fig. 4C-D). The stage VI PD group, though, was easily distinguishable from both controls (p < 0.001) and stage IV PD (p = 0.011 and p < 0.001), based on either deposit particles (Fig. 4C) or LBs (Fig. 4D). Mean and SEM values depicted in Fig. 4B-D can be found in Suppl. Table 2. In stage VI, it was evident that the LBs were selectively accumu-lated in the deep layers of the ACC (p < 0.001), as has previously been reported ^11,17,84^. We did not, however, detect any differences in PLA distribution between the superficial (SGM) and deep (DGM) layers of the ACC, nor any interaction between group and region.

Further investigation of the stage VI PD group – in which there were substantial amounts of PLA signal, IHC deposit particles, and LBs – revealed an inverse logarithmic correlation between PLA particle density and IHC deposit particle density (Fig. 4F). In addition, we detected a positive logarithmic correlation between IHC deposit particle and LB density (Fig. 4G), but no direct correlation between PLA signal and LB density (data not shown). Thus, the patients with the lowest density of PLA particles were the ones in which the highest density of IHC deposit particles was found, in contrast to the findings by Roberts et al. in the cingulate cortex ^45^. This suggests that the MJF-14 PLA signal indeed represents early α-synuclein pathology, which is sequestered into early types of deposits detected by IHC and later matures into LBs. To test this hypothesis, we revisited some of the DLB tissue stained for both MJF-14 PLA and pS129 (Fig. 3) and did a direct comparison of PLA signal in LB-positive and neighbouring LB-negative large neurons. Intriguingly, the average PLA area was 36% higher in LB-negative than LB-positive large neurons, supporting a process where oligomers are sequestered into inclusions (Suppl. Fig. 4).

We also looked further into the distribution of PLA signal between neuronal cell bodies and the neuropil (see Fig. 4H and methods for segmentation definition). Overall, there was a strong linear correlation between neuronal and neuropil PLA with R = 0.938 (p < 0.001), and around 30% of the total PLA particles were localized in neuronal cell bodies, i.e., in nuclei or surrounding cytoplasm (33.6% for PD stage IV and 28.9% for PD stage VI, Suppl. Table 4). Thus, the majority of the PLA signal was located away from the neuronal nuclei, in what is here denoted neuropil but also comprises axons, dendrites, and extra-neuronal matrix including various glial cells. A prime candidate for its principal location is the presynaptic and axonal compartments, which are believed to be the earliest affected regions of the neurons ^81,85–88^. Looking to the neuronal cell bodies, 6.76% of the neurons in controls contained PLA signal, while this proportion was almost 10-fold higher in PD, with 70.5% in stage IV PD and 63.5% in stage VI PD but with considerable intra-group variation (Fig. 4E and Suppl. Table 3). Additionally, the average load of PLA in affected neurons was also increased in the PD groups, both in deep and superficial grey matter (Fig. 4I-J and Suppl. Table 4). The proportion of PLA-containing neurons was also much higher than the corresponding neuronal proportions containing deposit particles in general (approx. 4.7% in stage IV and 10.7% in stage VI PD) or LBs specifically (4.13% in stage VI PD; Suppl. Table 3). There was no significant difference in the number of neurons per region of interest between the three groups (p = 0.103).

To appraise whether the PLA particles accumulated in a specific part of the neurons, we did a semi-quantitative single-neuron analysis, assessing PLA distribution between nuclear and cytosolic compartments. This revealed that the majority of PLA-containing neurons in both PD groups had signal either solely in the cytoplasm or in both nucleus and cytoplasm (Fig. 4K, Suppl. Table 5). The mixed distribution (PLA particles in both nucleus and cytoplasm) was highly specific for PD and did not occur in controls. Comparing PLA staining in the PD cases (Fig. 4) with the DLB cases (Fig. 3) led us to note that while nuclear PLA particles were found in both PD and control cases with chromogenic PLA, the fluorescent PLA appeared to mainly detect cytoplasmic and neuropil PLA in the DLB cases (Fig. 3C vs. Fig. 4A). This distinction appears to be methodology-related (i.e., the chromogenic setup may be more sensitive than the applied fluorescent PLA channel) rather than disease-related, as the same DLB cases stained with chromogenic PLA do display nuclear PLA particles (Suppl. Fig. 5A, top).

Like for the total analysis of PLA (Fig. 4B), we did not find any obvious differences in the PLA distribution (nuclear vs. cytosolic, Fig. 4J) between neurons in stage IV and VI PD in the ACC, perhaps due to the variation in PLA-detected pathology within each group. Still, our results underscore the potential of single-neuron PLA analysis as a supplement to the previously seen semi-quantitative general tissue scoring methods.

### A fraction of neuropil MJF-14 PLA signal localizes to the presynaptic terminal

Following our observations on the PD cohort in the anterior cingulate cortex, leading to the realization that the majority of PLA signal is located in the neuropil, we wanted to explore the exact localization of these neuropil PLA signals. As the structures in the neuropil and the PLA signals themselves are quite small, the diffraction limit significantly restricts our ability to do proper co-localization analysis – unless a super-resolution approach is used. Therefore, we set out to combine our PLA protocol with ONE microscopy to push the resolution as far as possible (Fig. 5A). Since the choice of fluorophores is crucial for good results with expansion, we switched to using PLA probes and kits from Navinci, where we acquired a custom-made kit that allowed labelling with multiple different fluorophores depending on our need.

**Figure 5:**
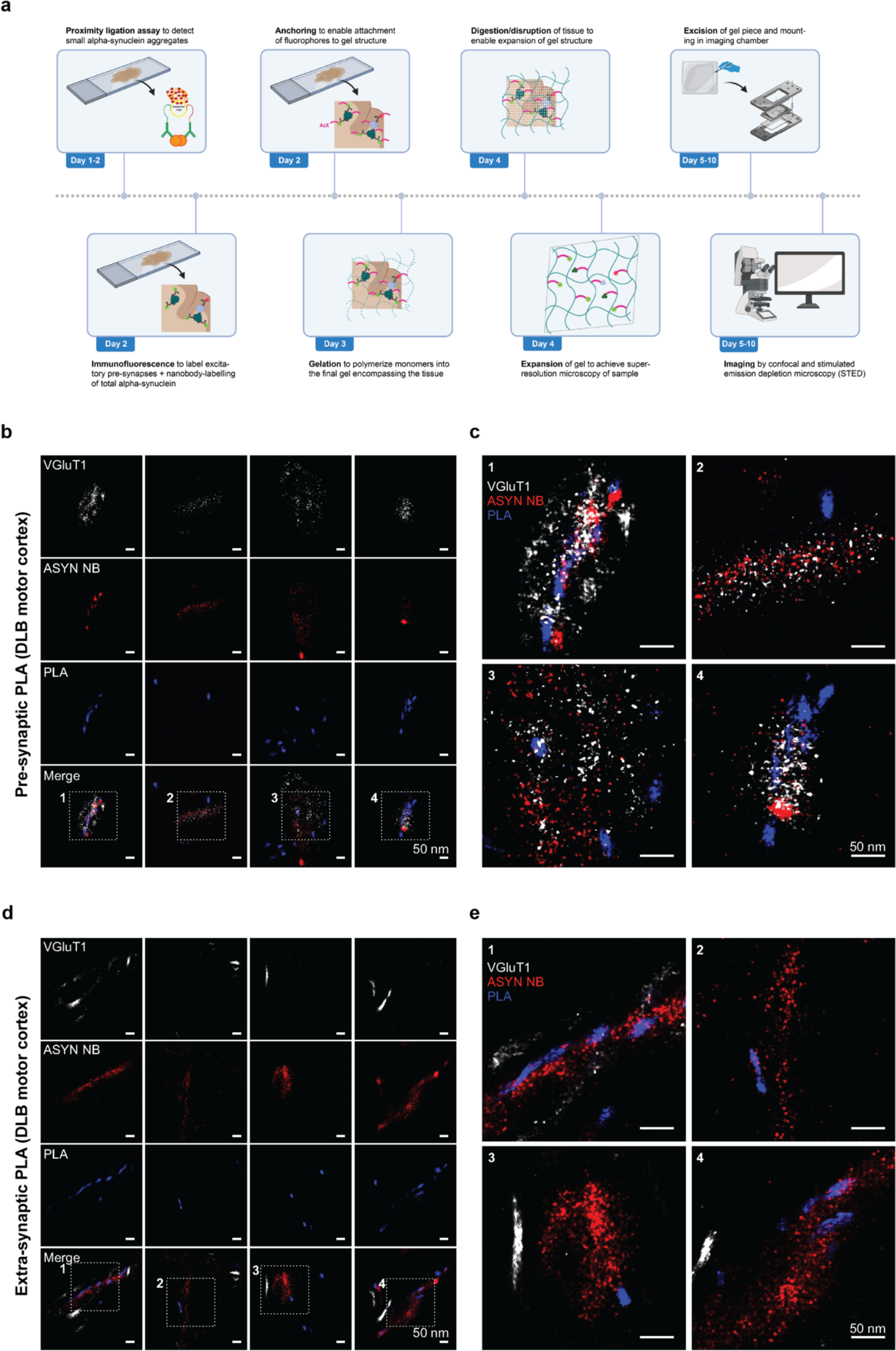
Applying ONE microscopy to PLA imaging decisively identifies α-synuclein aggregates at the presynaptic terminal in DLB motor cortex. **A.** Experimental setup of ONE micrscopy combined with MJF-14 PLA. Samples were stained for MJF-14 PLA, then VGLUT1 and total α-synuclein, before tissue expansion and ONE imaging of gels. **B.** Individual channel + merged images with four examples of presynaptic PLA signal in DLB motor cortex. VGLUT1 (white) labels excitatory presynapses, MJF-14 PLA (blue) labels small α-synuclein aggregates, and Asyn-Nb (red) labels total α-synuclein. **C.** Magnified panels 1-4 from **B** showing co-localization of PLA signal with both total α- synuclein and VGLUT1. **D.** Individual channel + merged images with four examples of extra-synaptic PLA signal in DLB motor cortex. VGLUT1 (white) labels excitatory presynapses, MJF-14 PLA (blue) labels small α-synuclein aggre-gates, and Asyn-Nb (red) labels total α-synuclein. **E.** Magnified panels 1-4 from **D** showing co-localization of PLA signal with total α-synuclein but not with VGLUT1. All scale bars = 50 nm. Panel A is created with BioRender.com.

We chose to study motor cortex from one of our DLB cases and compared it with a non-neurodegenerative control. Initial chromogenic PLA staining was performed to i) compare the MJF-14 PLA from Duolink and Navinci and ii) to form an impression of the PLA signal distribution in these cases for the following expansion experiments. Overall, MJF-14 PLA staining using kits from Duolink and Navinci resulted in similar staining patterns, with some parts of the DLB motor cortex mainly containing neuropil PLA and others mainly in the cell body, both nuclear and perinuclear (Suppl. Fig. 5A, top). The non-neurodegenerative control motor cortex was virtually blank with both company kits (Suppl. Fig. 5A, bottom). On closer inspection, we observed a strong labelling of what appeared to be LBs/inclusions with the Navinci MJF-14 PLA, which were not found with the Duolink MJF-14 PLA (Suppl. Fig. 5A). This indicates that not only epitope presentation but also the company-specific proximity probes play a major role in determining which α-synuclein species can be recognized by the PLA.

Having confirmed that the MJF-14 PLA using Navinci probes and kits achieved a similar detection of signal in both neuropil and perinuclear/cell body compartments as the Duolink MJF-14 PLA, we continued with setting up the expansion PLA with this kit. We hypothesized that some of the neuropil PLA was likely located in the presynaptic terminal – the compartment where native α-synuclein primarily resides and the putative origin of α-synuclein aggregate pathology. Therefore, samples were labelled with fluorescent Navinci MJF-14 PLA, followed by immunostaining for excitatory presynapses (VGLUT1) and total α-synuclein using an α-synuclein-specific nanobody (Asyn-Nb) ^62^. Confocal imaging of non-expanded samples confirmed staining protocol functionality and did indeed indicate the likely co-localization of a fraction of the MJF-14 PLA signal with VGLUT1 in the DLB motor cortex (Suppl. Fig. 5B, arrows). As for the chromogenic application, virtually no PLA signal was found in the control motor cortex (Suppl. Fig. 5B).

Following the expansion procedure, gel pieces were excised and placed in a specially designed imaging chamber (see sketch in Fig. 5A), removing as much liquid from the gel as possible to allow stable STED imaging. From expanded samples, MJF-14 PLA signal was indeed found in small areas positive for VGLUT1, resembling the putative sizes of individual excitatory presynapses ^89–92^, evidently containing small α-synuclein aggregates (Fig. 5B-C). Examples of MJF-14 PLA signals not co-localizing with VGLUT1 were also found (Fig. 5D-E), while MJF-14 PLA signal generally coincided with total α-synuclein nanobody staining (Fig. 5B-E), confirming the presence of α-synuclein at sites of PLA signal. The usage of the α-synuclein nanobody also facilitated the precisely localization of the α-synuclein in our samples, as linkage errors become a significant factor to account for after the expansion procedure. Since the expansion procedure isotropically expands everything in our samples, linkage error, i.e. the distance between fluorophores (which we observe) and actual targets (which we want to locate) is also expanded. The size of the linkage error thus depends on the combined size of our antibody/nanobody and fluorophore. As such, a linkage error of at least 20 nm is expected for VGLUT1 signals (stained with primary + secondary antibody, approx. 150 kDa each) and for the PLA, a linkage error around 50-60 nm is not unexpected ^93^. In contrast, using tagged nanobodies the typical linkage error is much lower, around 7-9 nm (approx. 15-20 kDa), allowing us to confidently localize the α-synuclein. Despite the limitations that the linkage errors pose in our experiments, the expansion PLA provides convincing documentation of α-synuclein ag-gregates in excitatory presynapses.

### MJF-14 PLA does not work in mouse models

To assess the appearance and distribution of MJF-14 PLA in a mouse model of PD, we initially compared signals in male human α-synuclein transgenic mice (Thy1-hasyn-Tg, line 61; ASO) and α-synuclein knock-out (C57BL/6N-Snca^tm1Mjff^; ASKO) mice. Surprisingly, substantial amounts of MJF-14 signal were detected in both ASO and ASKO mice, with region-dependent variations, i.e., highest signal in midbrain and lowest in hippocampus (Fig. 6A). Technical negative controls, omitting either the PLA probe-conjugated antibodies or the ligase in the PLA reaction showed essentially no signal (Suppl. Fig. 1A), demonstrating that the signal was not due to incomplete blocking of endogenous peroxidases. Moreover, the signal was present in ASKO mice through different batches of antibodies/PLA probes (data not shown). We also compared the results of the MJF-14-conjugated Duolink probes (Sigma) with MJF-14-conjugated Naveni PLA probes (Navinci), which yielded a similar PLA signal in ASKO mice (Fig. 6A vs. B). It was not possible to dilute the PLA probes so that no signal was present in ASKO without simultaneously losing all signals in the ASO mice (data not shown).

**Figure 6:**
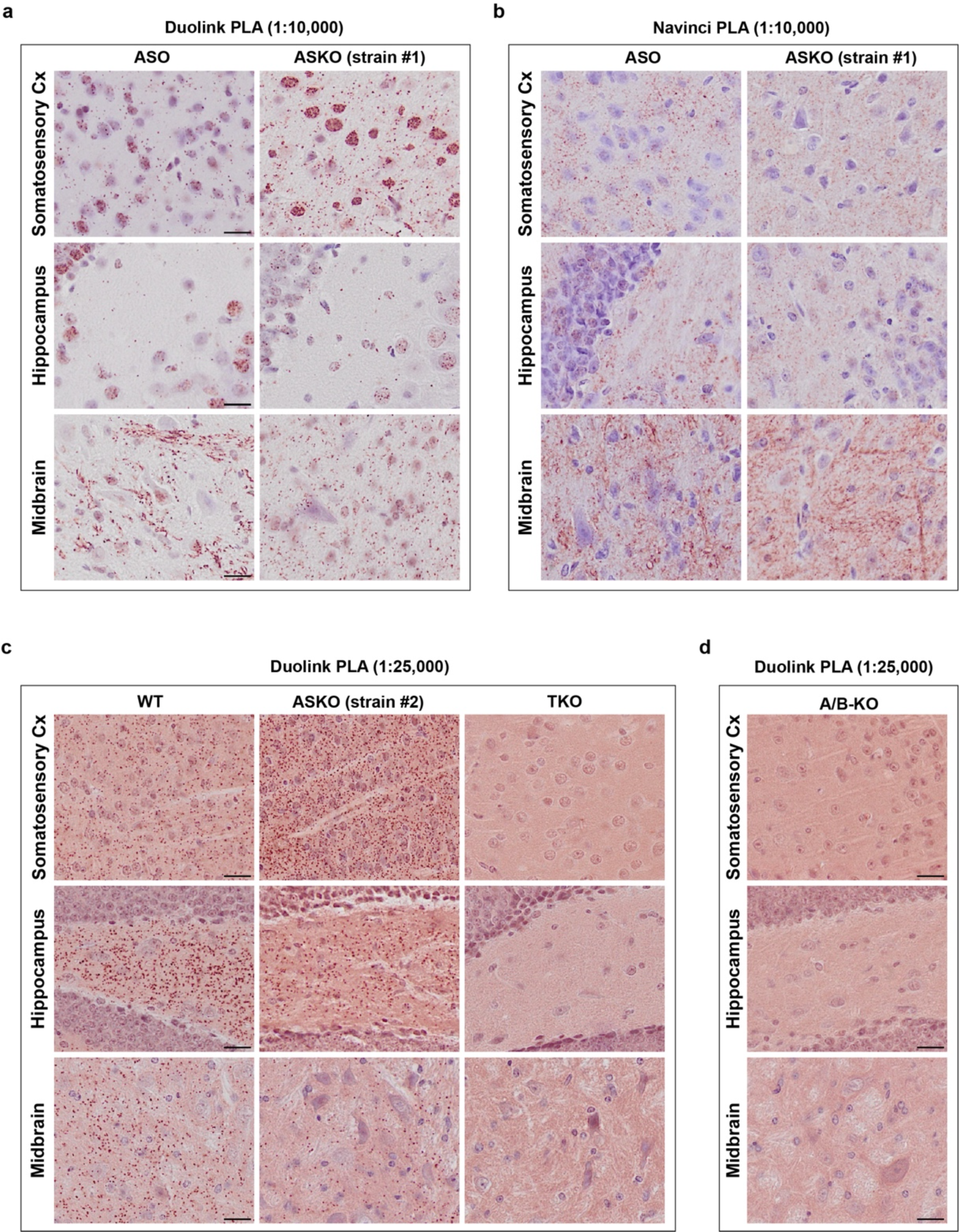
MJF-14 PLA signal is not specific for α-synuclein in mouse models. **A.** Comparison of MJF-14 PLA signal in α-synuclein transgenic (ASO) and α-synuclein knockout (ASKO strain #1) mice, using PLA probes and detection kits from Duolink (Sigma). **B.** Similar comparison of MJF-14 PLA signal as in **A**, but using PLA probes and detection kits from Navinci. **C.** MJF-14 PLA signal in a different α-synuclein knockout strain (ASKO strain #2), compared with WT, and triple α-/β-/γ-synuclein knockout mice (TKO). **D.** MJF-14 PLA signal in α-/β-synuclein double knockout mice (A/B-KO). All scale bars = 20 µm.

To exclude the possibility that the presence of the signal was caused by an artefact with our ASKO mouse strain (ASKO #1; C57BL/6N-Snca^tm1Mjff^), we obtained a second ASKO mouse strain (ASKO strain #2; B6(Cg)-*Snca*^tm1.2Vlb^/J) for comparison. This second ASKO strain also stained positive for MJF-14 PLA, albeit with more signal in the cortex and hippocampal regions and less in the midbrain, demonstrating that MJF-14 PLA positivity in ASKO mice is consistent across strains (Fig. 6C). To examine whether the signal could be due to binding to another mouse synuclein isoform, we also included α-/β-/γ-synuclein triple knockout (TKO) and α-/β-synuclein double knockout mice (A/B-KO) in the analysis. The TKO mice were completely negative for MJF-14 PLA (Fig. 6C), indicating a cross-detection of either β- or γ-synuclein, most likely β-synuclein where the 5 C-terminal residues are identical to α-synuclein ^94^. Indeed, MJF-14 PLA-staining of A/B-KO mice also didn’t produce any signal, determining mouse β-synuclein as the culprit (Fig. 2D). MJF-14 immunohistochemistry on parallel sections also resulted in staining in both WT and ASKO #2 mice, while TKO and A/B-KO mice remained blank, confirming that the signal presence in ASKO mice is antibody-dependent (Suppl. Fig. 1B). We, therefore, concluded that the MJF-14 antibody is not suitable for PLA in mouse models due to the strong background staining in ASKO mice, which could not be separated from potentially true α-synuclein aggregate staining in transgenic mice.

## Discussion

In this study, we present a novel α-synuclein PLA based on the aggregate-specific MJF-14 antibody and evaluate its functionality in synucleinopathy models as well as post-mortem patient tissue. We show that aggregation is a prerequisite for MJF-14 PLA signal formation, based on the significant decrease in signal in our cell model upon treatment with aggregation inhibitor ASI1D. No similar decrease in syn211 PLA signal was seen following ASI1D-treatment, despite the fact that Roberts et al. also showed a need for multimerization, based on bimolecular fluorescence complementation, in order to generate syn211 PLA signal ^45^. The differences between the MJF-14 and the syn211 PLAs observed in this study may be caused by the syn211 PLA targeting multimers – including potential physiological polymers ^75–78^. As reviewed by Estaun-Panzano et al., it is important to remember that the PLA technique in itself does not provide any selectivity towards pathological aggregates ^95^, but the specificity of the MJF-14 antibody for pathological aggregates may confer such quality.

The utility of the MJF-14 antibody, in both PLA and IF applications, is further underscored by its potential in models combining it with MJF-14-invisible stealth PFFs to induce α-synuclein pathology. This allows a confident examination of endogenous α-synuclein aggregation without the risk of cross-detection of the exogenous PFFs. As the stealth PFFs were also engineered to be S129A-mutated, and thereby non-phosphorylatable at serine-129, pS129-α-synuclein-specific antibodies will also only stain endogenous α- synuclein in this setup ^48^. Thereby, these tools enable detailed studies into the early aggregation and phosphorylation processes and can be combined with a range of model systems.

The clinical material studied in this paper consisted of a PD/control cohort from anterior cingulate cortex (n = 10 per group) and a DLB/control cohort from the superior frontal cortex and motor cortex (n = 2 per group). The PD cohort was used to compare MJF-14 PLA directly with IHC and study the distinction in pathology at stage IV vs. VI PD, while the DLB cohort was used to study the LB-labelling capacity of the MJF-14 PLA and compare its detection ability between different company kits (Duolink vs. Navinci MJF-14 PLA). Although the clinical material was rather limited in this study, a cohort with 10 stage IV and 10 stage VI PD patients is larger than what has previously been published in PLA studies of α-synuclein pathology and does bring forth new clinical insight as discussed below. In both the PD and DLB cohorts, we found striking amounts of MJF-14 PLA signal, indicating a much more widespread dysfunction in these tissues than what is revealed using traditional IHC staining techniques with either pS129- or confor-mation-specific α-synuclein antibodies. Furthermore, the comparison of stage IV and VI PD implied the ability of the MJF-14 PLA to detect pathology that markedly precedes bona fide inclusions in human brains as stage IV is defined by being essentially devoid of LBs in the ACC. Thereby, our data support the findings from Sekiya et al., who demonstrated oligomeric pathology using the syn211 PLA in multiple hippocampal and cortical regions of stage III-IV PD without concurrent presence of Lewy-related pathology ^74^. Although they did not examine the ACC, a previous paper from the same group demonstrated PLA-positivity in 4/5 PD patients in the ACC, in the form of “punctate” staining ^72^. While difficult to directly compare results between studies, all PD cases in our cohort were PLA-positive in the ACC. This difference could be explained by a higher sensitivity of the MJF-14 PLA or by the detection of different oligomers/polymers by the two PLAs, as mentioned above. Alternatively, the use of distinct Braak stages could influence the findings, as Sekiya et al. included 5 cases of stage III-VI, of which one stage III case was void of PLA signal in the ACC ^72^, while we examined 10 stage IV and 10 stage VI cases.

In the PD cohort, we found considerable variability in the density of PLA pathology between cases in the PD groups – a characteristic that was independent of their Braak stage. Half of the patients in both stage IV and VI in our cohort displayed PLA pathology in more than 80% of the neurons detected by automated analysis, suggesting an extensive neuronal dysfunction in this area. Conversely, in stage VI, there were 3/10 patients in which fewer than 25% of the neurons (range 11.7 – 22.9%) contained PLA pathology (see Fig. 4E). This suggests that the Braak staging is likely not sufficient for properly characterizing the molecular α-synuclein pathology in PD patients, as has previously been indicated ^20,96–98^, but should be complemented by other methods to increase our understanding of disease including examination of various post-translational modifications ^99^. Whether this inter-individual variation seen in the two PD groups might be caused by different subtypes of PD, for instance brain-first and body-first as recently suggested by Borghammer and colleagues ^100–103^, or related to e.g. genetic autophagy-lysosomal-related risk factors of sporadic PD, can at present only be speculated upon. Nevertheless, this degree of variability between patients of the same Braak stage exposed by proximity ligation has not previously been re-ported – perhaps because earlier studies have only used smaller groups of mixed disease stages ^45,72,74^.

To characterize the α-synuclein pathology in the PD cohort, we employed three different measures: PLA particles, deposit particles (i.e., any accumulation of aggregated α-synuclein with a minimum length of 2 µm), and LBs (defined by two independent researchers), the latter two based on MJF-14 IHC. From this, we discovered an inverse correlation between PLA particle density and the density of α-synuclein deposit particles (detected by IHC) in the stage VI PD patients. Combined with the observation that LB-containing large neurons in DLB harbour less PLA signal than neighbouring LB-negative neurons, this indicates that α-synuclein inclusions are able to sequester PLA-positive oligomers and convert them into (Duolink MJF-14) PLA-negative species. As multiple studies have shown that the classical Lewy pathology is not a good measure for estimating either neuronal death or symptomatic burden of patients ^20,21,104^, our results raise the speculation whether the PLA pathology might correlate better with impact on cellular functions or disease progression. Additionally, our results further weigh into the discussion of whether the LB is in itself a toxic species, a – perhaps failing – protective response that sequesters potentially toxic PLA-positive species, or simply a bystander in synucleinopathies ^11,18,19,21,22,104,105^.

A curious observation in this study was that not only the selected antibody but also the specific brand of PLA kit plays a crucial role in determining which α-synuclein species are detected in the PLA. As such, the Duolink MJF-14 PLA did not show efficient labelling of LBs in either DLB frontal and motor cortex or in PD anterior cingulate cortex. In contrast, the Navinci MJF-14 PLA, although only tested on the DLB motor cortex, displayed a strong labelling of LBs in addition to the labelling of non-inclusion aggregate pathology. As the specific kit formulations, including the structure of PLA probes, are proprietary to both Duolink and Navinci, we can only conclude that careful validation is necessary when switching between kit suppliers. Adding to the complexity, although Roberts et al. also showed an overall predilection of the syn211-PLA towards non-LB pathology, their assay showed a stronger detection of cortical LBs than with regular immunohistochemistry in both PD and DLB patients ^45^. Collectively, we are left with two versions of the new MJF-14 PLA: a version with a high specificity towards early α-synuclein pathology and only sparse labelling of inclusions (Duolink), and another version that appears to label all α-synuclein pathology, whether organized into inclusions or not (Navinci). The discrepancy between the detected species with the two PLA kits is likely caused by distinct oligonucleotide linker composition, which could allow for more or less flexibility in epitope binding and subsequent linker ligation. For some applications, such as pathological examinations, the Navinci MJF-14 PLA could be ideal in that it would allow the examination of a wider spectrum of α-synuclein pathology in one slide. This would of course need validation to elucidate whether this PLA does indeed recognize LBs and other α-synuclein inclusions as efficiently as the current standard IHC methods. On the other hand, one could also imagine applications where a preference for non-inclusion α-synuclein pathology is more suitable, in which case the Duolink MJF-14 PLA would be superior. At the structural level, the different specificity of the two kits suggests that modifying the oligonucleotide linkers, which are conjugated to the MJF-14 antibody, holds promise for development of aggregate-or even strain-specific PLA reagents.

In this study, we present an assortment of analyses on PLA-stained samples, going beyond the scope of semi-quantitative measures based on scoring plates. These ranged from simple quantifications of PLA particle counts normalized to either cell number, field of view, or tissue area to more advanced quantifications of e.g. PLA in various neuronal compartments or PLA in neurons with/without LBs. An interesting observation, based on the fluorescent MJF-14 PLA on both human cortical neurons and neuroblastoma cells, was that not only PLA particle count but also intensity varied across conditions. This phenomenon is most likely caused by the presence of multiple rolling circle amplification products on the same aggregate, indicating that the PLA-positive aggregates detected may also differ in size between conditions. It opens for considering PLA signal as not simply a dot or particle of inconsequential intensity, but something where analysis of integrated density (PLA area times intensity) may give further insight into aggregate dynamics. While several protocols for PLA analysis have been published, none has suggested integrated density as a measure of PLA signal, perhaps because PLAs are mostly applied to study protein interactions rather than aggregate dynamics ^106–109^.

As previously outlined, this study is somewhat limited by low number of cases in several analyses, leaving many of the conclusions to be validated in larger studies. Nevertheless, our main purpose was to highlight the potential of proximity ligation assays in synucleinopathy research, particularly combined with other methods (such as super-resolution microscopy) and analyses (including more quantitative single-neuron approaches) than previously presented. For instance, by combining our PLA with expansion and STED microscopy, we demonstrated the presence of PLA-positive aggregates in excitatory pre-synapses in DLB motor cortex. Here, further studies to follow up on the other cellular compartments containing PLA signal as well as their quantitative distribution during the course of disease could be of great value. Additional studies of multiple brain regions for their PLA-detected α-synuclein pathology are also highly motivated, especially regions previously considered negative for pathology, such as the cerebellum in PD, where PLA signal has already been demonstrated in a few cases ^72^.

Other limitations of this study concern the segmentation of signals in the PD cohort, in particular the definition of neuronal nuclei and their respective cytoplasm. Since the sections included in the study were only counterstained with the nuclear dye haematoxylin, neuronal nuclei were distinguished from glial nuclei solely based on differences in size, which does not allow a clear-cut distinction. Similarly, cytoplasm was only rarely stained by the counterstain and, consequently, in the analysis the cytoplasm was estimated as a circular border around the nucleus, based on average measurements of cytoplasm within neurons with visible cytoplasm. These limitations could be overcome by the use of other counter-staining methods, or co-staining with specific neuronal or other cell type markers in future. In the automation process, we were additionally challenged by variations in the counterstaining efficiency, which left some of the sections rather faintly counterstained, despite being stained in the same batch as other sections with much brighter counterstain. In the end, we had to divide sections into two groups depending on counterstain efficiency and develop distinct classifiers for these groups. Although not optimal, it was not possible to achieve anything close to a satisfactory segmentation using only one classifier, while using two classifiers, we achieved a sensitivity for recognition of neuronal nuclei of 85% (Suppl. Fig. 1).

Also, as there was no significant difference between controls and the two PD groups in counterstain efficiency (p = 0.37), we were assured that the classifiers did not accidentally skew the analysis.

Lastly, we discovered that the MJF-14 PLA was not functional in mouse models due to a cross-detection of β-synuclein and a resulting large background signal, even in α-synuclein knockout mice. This cross-detection did not appear to be an issue for human tissue, as demonstrated by the blank regions of interest in control ACC (Fig. 4). Nevertheless, this means that there are currently no published options for α- synuclein PLA on mouse models only expressing mouse α-synuclein, as the syn211 antibody is human-specific and therefore cannot recognize aggregates of mouse α-synuclein.

In conclusion, we here demonstrate a novel α-synuclein PLA with increased specificity towards early oligomeric α-synuclein pathology and highlight novel combinations with super-resolution imaging techniques and with specialized stealth PFFs, which are invisible to the MJF-14 antibody in both PLA and IF applications. We underscore the fact that α-synuclein pathology detected by MJF-14 PLA in both PD and DLB precedes and is much more widespread than Lewy pathology detected by conventional IHC, with on average two-thirds of the neurons affected by PLA pathology in the cingulate cortex of both stage IV and VI PD patients. In comparison, only 4% of neurons contained LBs even in stage VI patients. In addition, we provide a method for extensive automated analysis of chromogenic PLA-stained tissue sections, thereby setting the scene for comprehensive large-scale clinical studies of non-inclusion α-synuclein aggregate pathology, which has not hitherto been appreciated.

