## Supplementary figures and tables for "MJF-14 proximity ligation assay detects early non-inclusion alpha-synuclein pathology with enhanced specificity and sensitivity"

**Supplementary figure 1**

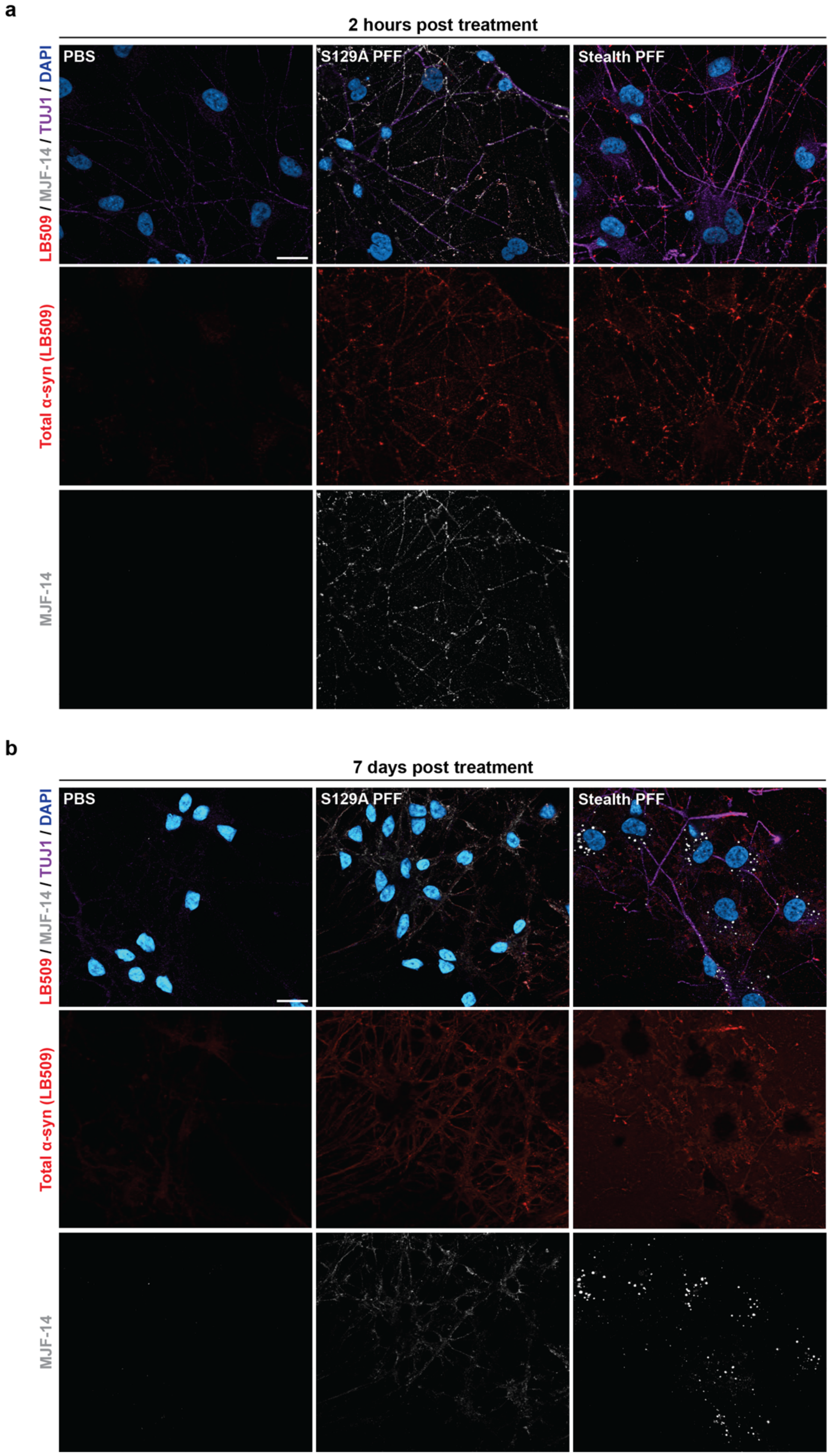


**Supplementary figure 1: Stealth PFFs are invisible to the MJF-14 antibody and increase signal-to-noise ratio in immuno-fluorescence staining of human cortical neurons. A-B.** Immunofluorescence of human cortical neurons treated with either S129A PFFs, stealth PFFs (S129A-α-synuclein-141G), or PBS, and fixed after 2 hours (**A**) or 7 days (**B**). Representative images of cultures immunostained with MJF-14 (grey), total α-synuclein (LB509, red), βIII-tubulin (purple), and DAPI nuclear stain (blue) show strong detection of exogenous S129A PFFs with MJF-14 after 2 hours, while stealth PFFs don't generate any signal (**A**). After 7 days, increased MJF-14 staining is seen with both S129A and stealth PFF treatment, but considerable background staining, perhaps from non-internalized PFFs, is still present in S129A PFF cultures. In contrast, α-synuclein deposits in stealth PFF cultures are easily identifiable based on the MJF-14 staining (**B**). Scale bars = 20 µm.

### Supplementary figure 2

**a**

Group 1: Intense neuronal counterstain

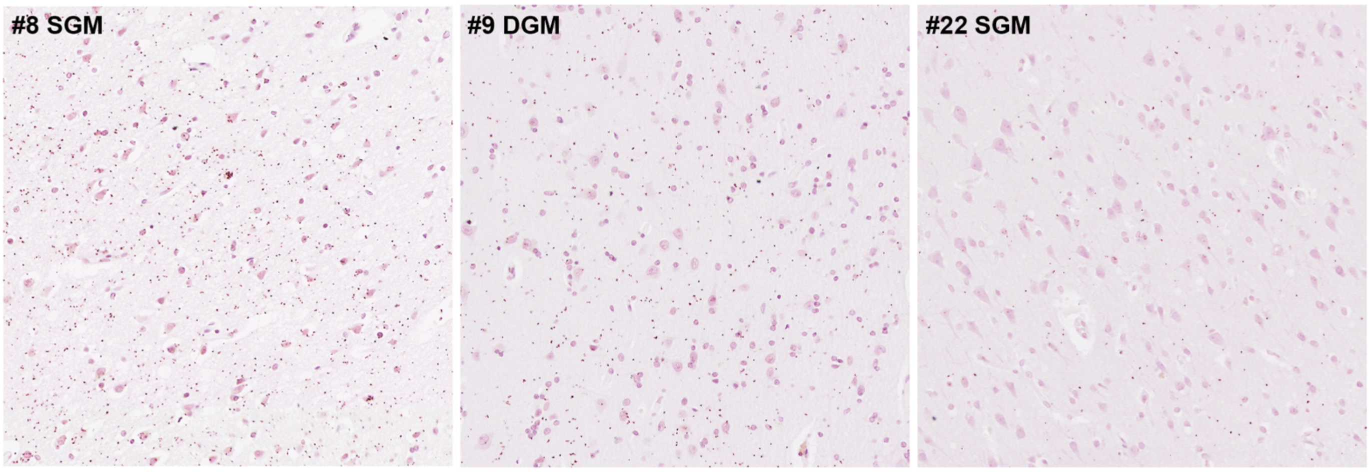


**b**

Group 2: Weak neuronal counterstain

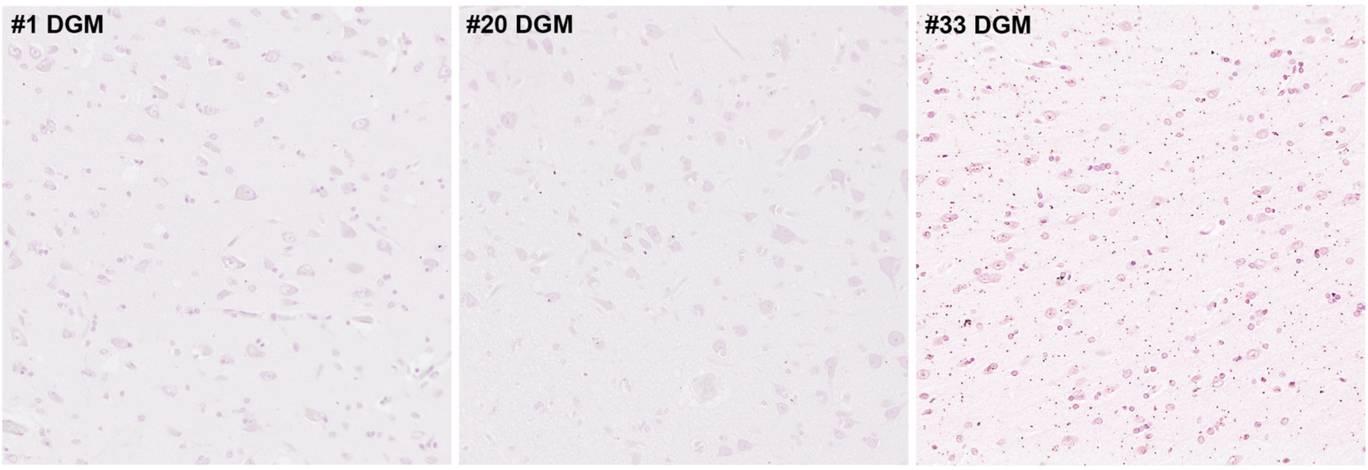


**c**

| | Sensitivity | Precision | Accuracy |
|---|---|---|---|
| **Definition** | $\frac{true\ positive}{true\ positive\ +\ false\ negative}$ | $\frac{true\ positive}{true\ positive\ +\ false\ positive}$ | $\frac{true\ positive}{true\ positive\ +\ false\ positive + false\ negative}$ |

**d**

Group 1: Intense counterstain

| | Sensitivity | Precision | Accuracy |
|---|---|---|---|
| **#8 SGM** | 0.71 | 0.85 | 0.63 |
| **#9 DGM** | 0.91 | 0.69 | 0.65 |
| **#22 SGM** | 0.88 | 0.70 | 0.64 |
| **Mean** | 0.83 | 0.75 | 0.64 |

**e**

Group 2: Weak counterstain

| | Sensitivity | Precision | Accuracy |
|---|---|---|---|
| **#1 DGM** | 0.90 | 0.53 | 0.50 |
| **#20 DGM** | 0.98 | 0.68 | 0.67 |
| **#33 DGM** | 0.74 | 0.77 | 0.61 |
| **Mean** | 0.87 | 0.66 | 0.59 |

**Supplementary figure 2: Optimization of automated segmentation of chromogenic PLA-stained human anterior cingulate cortex. A-B.** ROIs used for optimization of PLA signal and nuclei segmentation grouped into either intense counterstain (**A**) or weak counterstain (**B**). Numbers indicate case ID, while SGM and DGM signify superficial and deep grey matter, respectively. The total number of PLA particles per image and classification into counterstain group for all cases can be found in Suppl. Table 1. **C.** Definition of parameters for evaluation of segmentation classifier performance. **D-E.** Final parameter values for intense (**D**) and weak (**E**) neuronal counterstain groups, respectively.

**Supplementary figure 3**

### Strategy for image segmentation and analysis

**1** Selection of test cohort for development of image classifier + manual annotation of test cohort

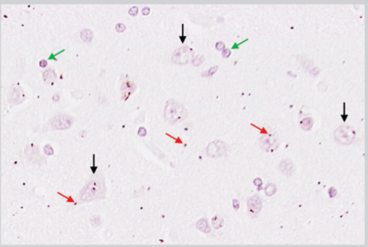


**2** Training and validation of classifier

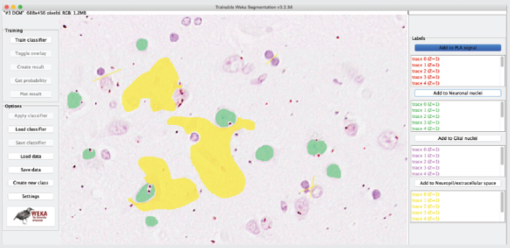

**3** Get probability maps for nuclei and PLA signal

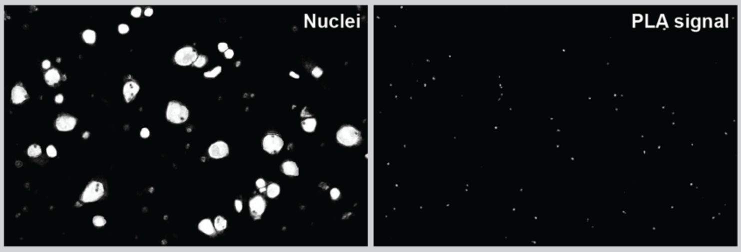


Add probability threshold
Filter on size (neurons >350 pixels)

Add probability threshold
Watershed to separate particles

**4** Analyze total, neuronal and extra-cellular/glial areas, PLA dots and neuronal count

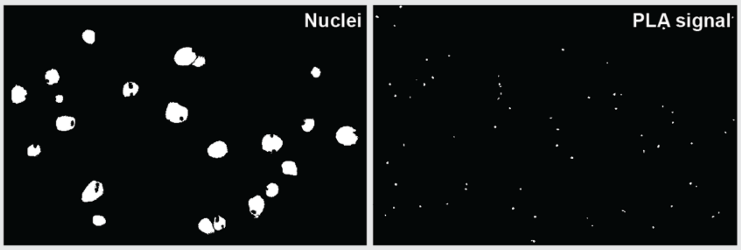


**5** Analyze each neuron individually

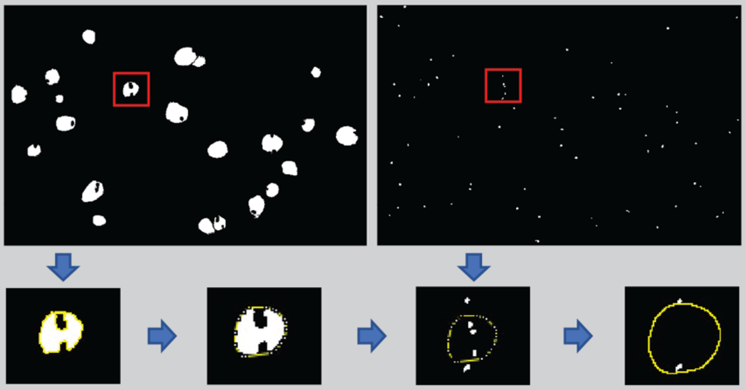

Select one neuronal nucleus → Fit convex hull (include indentations in nucleus) → Apply nuclear selection to PLA image, count dots and clear → Expand selection to include cytoplasm (3.5 μm border)

**6** Group neurons according to table

| No. of Neuron (Raw Counting data) | | | | | | | | |
|---|---|---|---|---|---|---|---|---|
| Nuclear type | | Cytoplasmic type | | Mixed type | | | | |
| 1 PLA dot in nucleus + 0 PLA in cell body | >1 PLA dot in nucleus + 0 PLA in cell body | 1 PLA dot in cell body + 0 PLA in nucleus | >1 PLA dot in cell body + 0 PLA in nucleus | 1 PLA dot in nucleus + 1 PLA dot in cell body | >1 PLA dot in nucleus + 1 PLA dot in cell body | 1 PLA dot in nucleus + >1 PLA dot in cell body | >1 PLA dot in nucleus + >1 PLA dot in cell body | No. of neurons without PLA signal |

**Supplementary figure 3: Strategy for image segmentation and quantification of PLA in human anterior cingulate cortex sections.** First, a test cohort consisting of 6 images was selected for the development and optimization of the automated classifier to segment the images (comprising differences in both PLA signal density and counterstain; see Suppl. Fig. 2). The test cohort was manually annotated before training of the classifier, which was then validated in its ability to correctly classify PLA signals and both glial and neuronal nuclei. For the classification of the entire ACC cohort, each image was then segmented and probability maps for nuclei and PLA signal were obtained. To define neuronal nuclei, a probability threshold was added, and particles larger than 350 pixels (36.7 $\mu m^2$, corresponding to an approx. diameter larger than 7 µm) were considered neuronal nuclei. To define PLA particles, a probability threshold was similarly added, fused particles separated using Watershed segmentation and particles with a size of 4-30 pixels (approx. 0.4-3.2 $\mu m^2$) counted. In addition, the total number of neuronal nuclei was counted, and total tissue area as well as glial/neuropil/extracellular area were computed, along with PLA particle counts in these compartments. For the single neuron analysis, each neuron was then analysed individually with 1) selection of nucleus, 2) fitting of a convex hull to include the indentations left by the PLA signal in the nuclei map, 3) application of the nuclear selection to the PLA signal map, counting of particles and clearing, and 4) expansion of nuclear selection by 3.5 µm to estimate cytoplasm, followed by counting of cytoplasmic PLA particles. Finally, each neuron was grouped in one of nine groups according to nuclear and cytoplasmic PLA particle counts.

### Supplementary figure 4

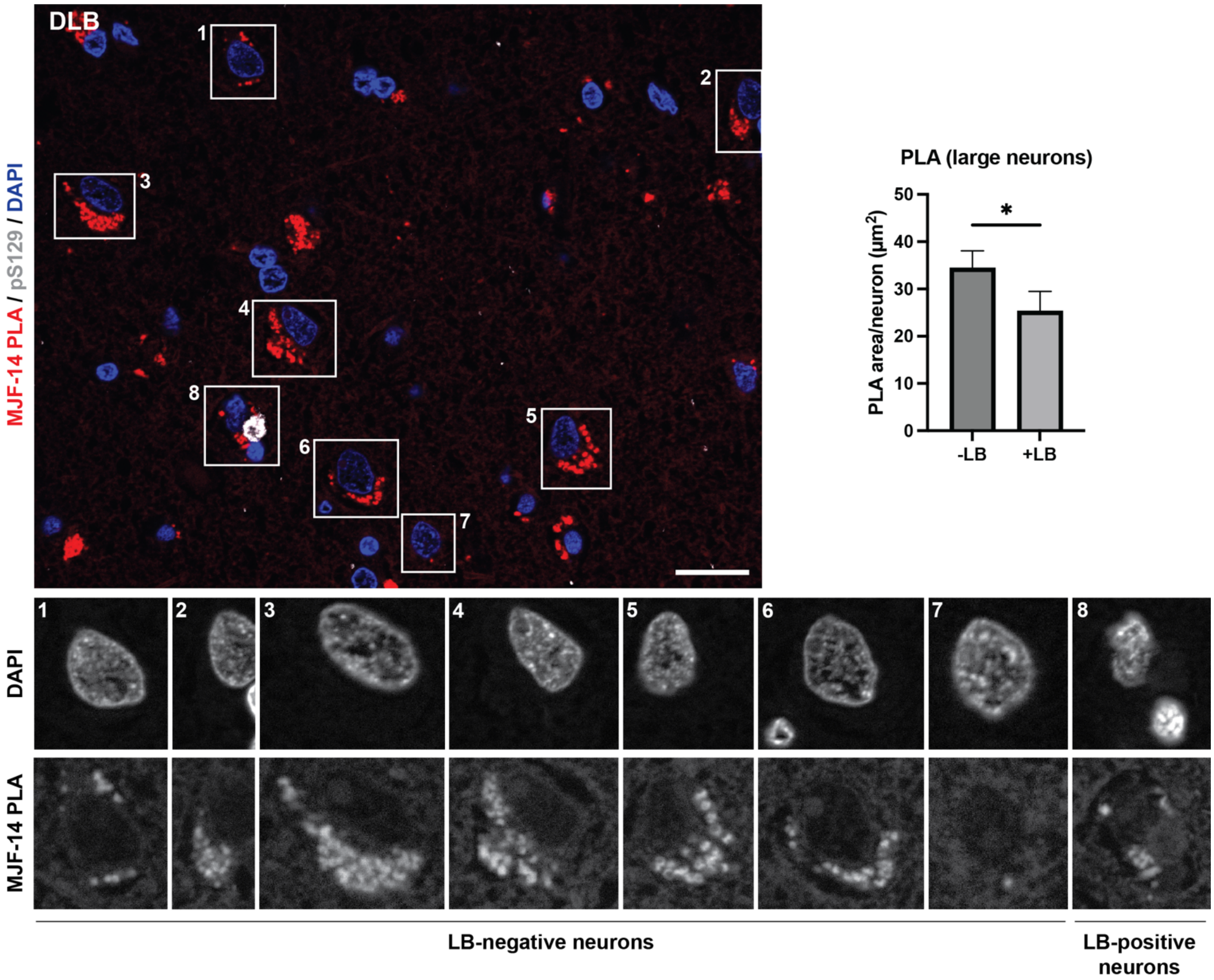


**Supplementary figure 4: LB-positive large neurons contain less PLA signal than their neighbouring LB-negative neurons.** Large neurons (outlined in white in merged channels image) from DLB frontal cortex samples were manually selected based on DAPI-staining, and single-neuron PLA images were extracted and grouped into LB-positive (neuron 8) and LB-negative (neurons 1-7) neurons. Merged image displays PLA signal in red, pS129-positive LBs in grey, and nuclei in blue, while single-neuron images are shown in greyscale. LB-positive neurons contained significantly less PLA signal than neighbouring LB-negative neurons ($p=0.0451$). Groups were compared using a Wilcoxon matched pairs signed rank test, as they did not pass normality. Scale bar = 20 µm. n=28 images were analysed (containing a total of 30 LB-positive neurons and 114 LB-negative large neurons). Data are displayed as mean ± SEM of PLA particle area/image. * $p<0.05$.

**Supplementary figure 5**

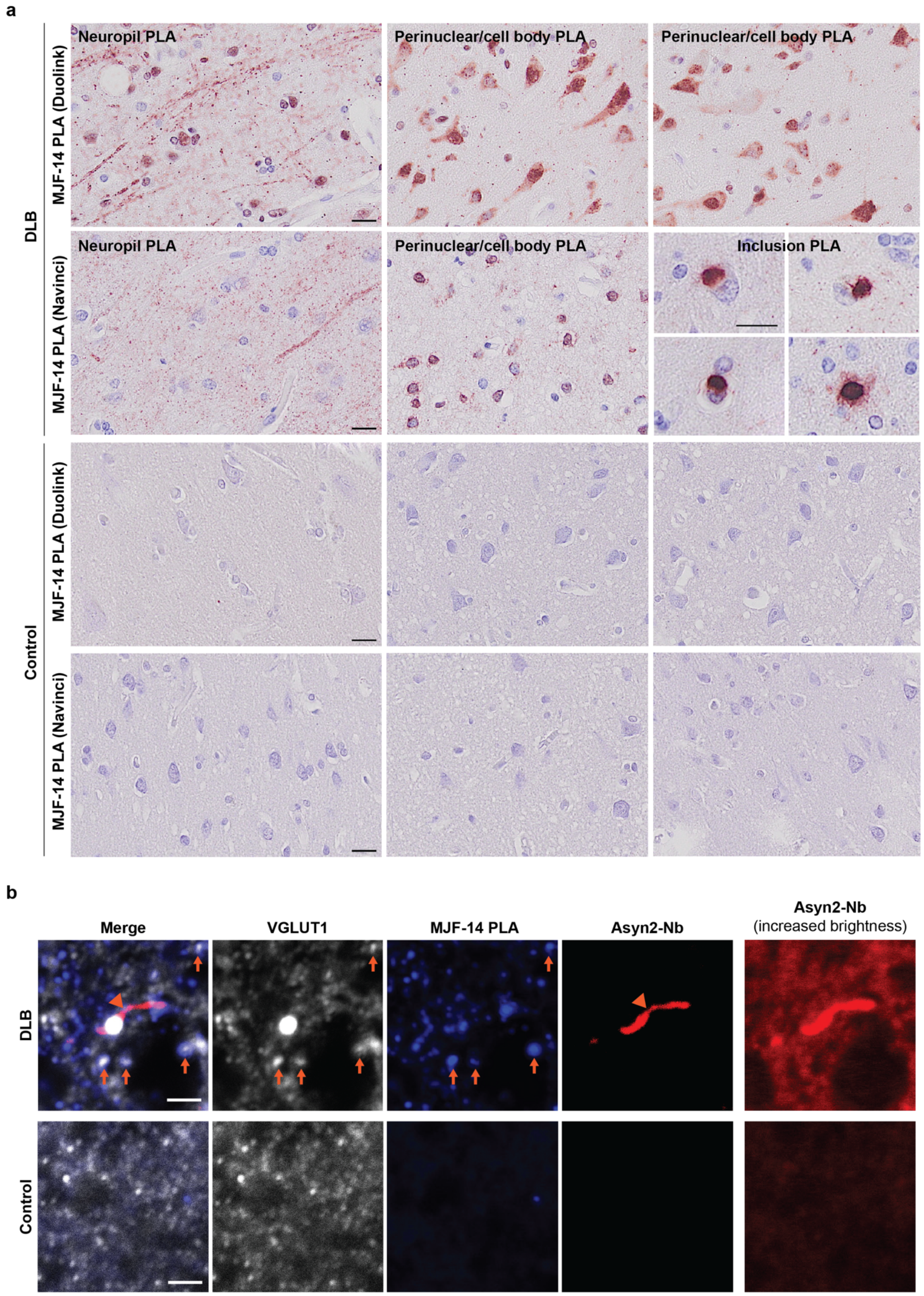


**Supplementary figure 5: Non-expanded supplementary stainings for expansion PLA experiment. A.** Chromogenic PLA overview of the motor cortex from DLB (top) and non-neurodegenerative control (bottom), stained with MJF-14 PLA from either Duolink or Navinci, exemplifying areas with preferential staining in the neuropil or the cell body. Note that strong labelling of inclusions is only seen with Navinci MJF-14 PLA. Scale bars = 20 µm. **B.** Confocal images from non-expanded staining samples in the expansion PLA setup, with excitatory presynaptic terminals (VGLUT1, white), $\alpha$-synuclein aggregate PLA (Navinci MJF-14 PLA, blue), and total $\alpha$-synuclein (Asyn2-Nb, red). A Lewy neurite (most likely located in an axon) is stained by Asyn2-Nb (arrowhead), while examples of apparent PLA-VGLUT1 co-localization are indicated by arrows. An increased brightness version of the Asyn2-Nb channel is shown on the right. Scale bar = 2 µm.

**Supplementary figure 6**

**a**

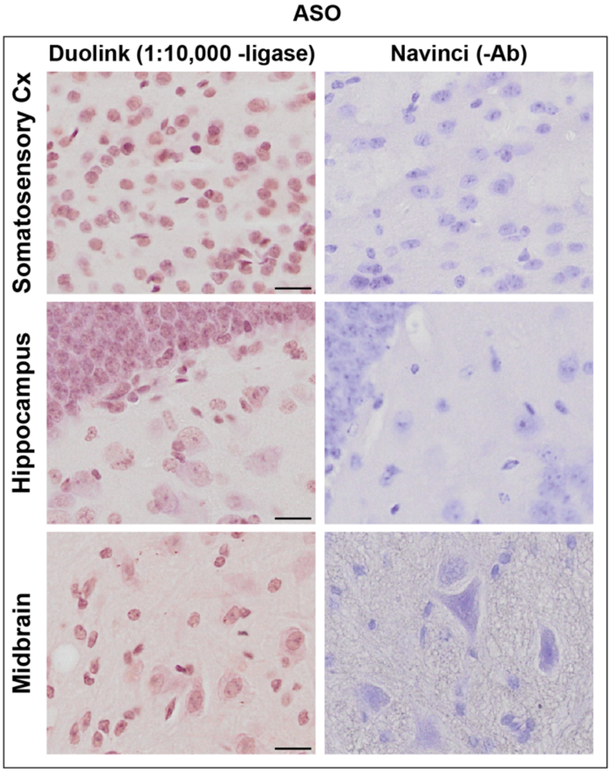


**b**

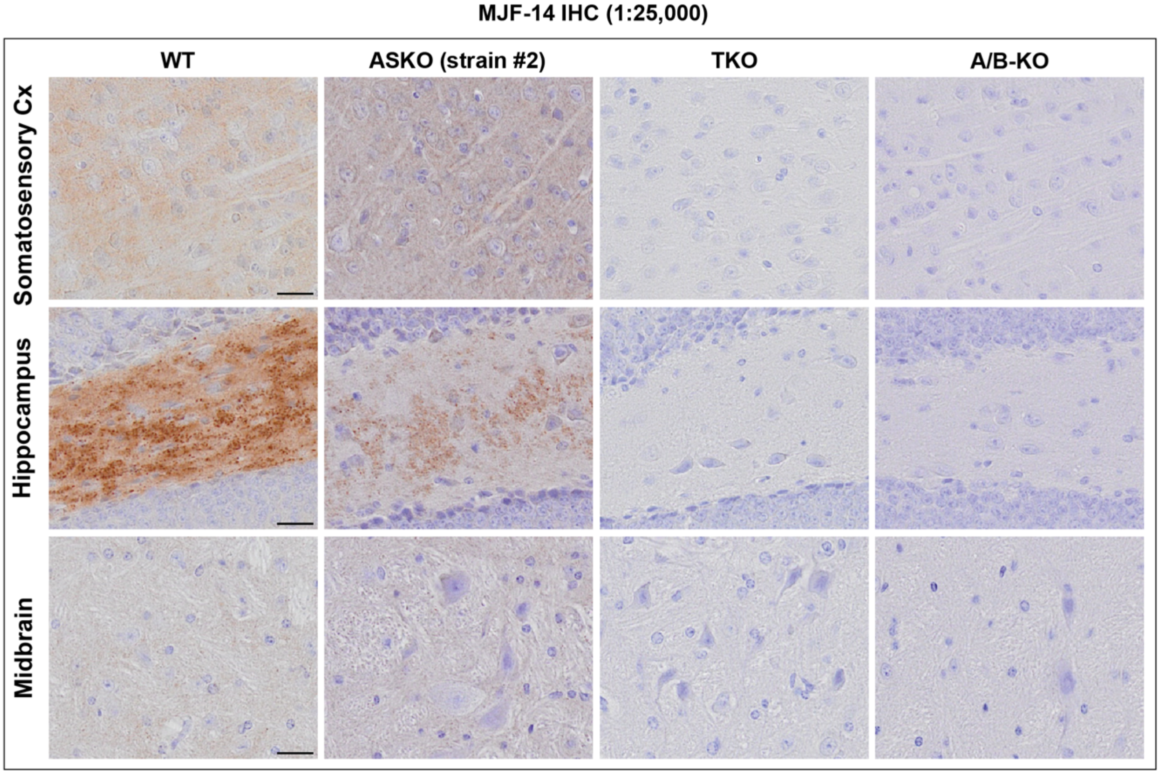


**Supplementary figure 6: Technical negative PLA controls and IHC staining of mouse models. A.** Technical negative control staining for MJF-14 PLA, lacking either the ligase in the PLA reaction (preventing signal formation) or the primary PLA antibody. The lack of signal in the controls demonstrate that insufficient blocking of endogenous peroxidase is not the cause for signal in ASKO mice. **B.** MJF-14 IHC also shows signal presence in ASKO mouse tissue, confirming the issue to be antibody-related. As both $\alpha$-/$\beta$-/$\gamma$-synuclein triple knockout and $\alpha$-/$\beta$-KO mice are blank, cross-detection of $\beta$-synuclein is most likely the cause of the signal. Scale bars = 20 µm.

### Supplementary tables

**Supplementary Table 1** PLA particles/case and division of cases by counterstain efficiency

| Section no. | Category | ACC region | Group 1: Intense counterstain | Group 2: Weak counterstain | PLA particles/image |
|---|---|---|---|---|---|
| 1 | Control | DGM | | X | 15 |
| | | SGM | | X | 4 |
| 2 | PD stage IV | DGM | | X | 349 |
| | | SGM | | X | 377 |
| 3 | PD stage VI | DGM | | X | 366 |
| | | SGM | | X | 212 |
| 8 | PD stage VI | DGM | X | | 896 |
| | | SGM | X | | 1413 |
| 9 | PD stage IV | DGM | X | | 574 |
| | | SGM | X | | 737 |
| 10 | Control | DGM | | X | 3 |
| | | SGM | | X | 3 |
| 11 | Control | DGM | | X | 17 |
| | | SGM | | X | 15 |
| 12 | PD stage IV | DGM | | X | 446 |
| | | SGM | | X | 64 |
| 13 | PD stage VI | DGM | | X | 105 |
| | | SGM | | X | 52 |
| 18 | PD stage IV | DGM | | X | 187 |
| | | SGM | | X | 237 |
| 19 | PD stage VI | DGM | X | | 736 |
| | | SGM | X | | 958 |
| 20 | Control | DGM | | X | 21 |
| | | SGM | | X | 34 |
| 21 | Control | DGM | | X | 5 |
| | | SGM | | X | 10 |
| 22 | PD stage IV | DGM | X | | 112 |
| | | SGM | X | | 192 |
| 23 | PD stage VI | DGM | X | | 60 |
| | | SGM | X | | 28 |
| 28 | PD stage VI | DGM | | X | 60 |
| | | SGM | | X | 110 |
| 29 | PD stage IV | DGM | X | | 1049 |
| | | SGM | X | | 1093 |
| 30 | Control | DGM | X | | 5 |
| | | SGM | X | | 22 |
| 31 | Control | DGM | | X | 37 |
| | | SGM | | X | 13 |
| 32 | PD stage IV | DGM | | X | 1031 |
| | | SGM | | X | 1109 |
| 33 | PD stage VI | DGM | | X | 949 |
| | | SGM | | X | 2771 |
| 38 | PD stage VI | DGM | | X | 933 |
| | | SGM | | X | 1389 |
| 39 | PD stage IV | DGM | | X | 1264 |
| | | SGM | | X | 881 |
| 40 | Control | DGM | | X | 4 |
| | | SGM | | X | 6 |
| 41 | Control | DGM | | X | 8 |
| | | SGM | | X | 18 |
| 42 | PD stage IV | DGM | X | | 441 |
| | | SGM | X | | 377 |
| 43 | PD stage VI | DGM | X | | 1006 |
| | | SGM | X | | 895 |
| 48 | PD stage VI | DGM | | X | 344 |
| | | SGM | | X | 1092 |
| 49 | PD stage IV | DGM | X | | 570 |
| | | SGM | X | | 536 |
| 50 | Control | DGM | X | | 238 |
| | | SGM | X | | 62 |

**Supplementary Table 2** Comparison of the signal density of PLA and IHC (data presented as mean ± SEM)

| | Controls | PD stage IV | PD stage VI |
|---|---|---|---|
| **Total PLA/$mm^2$** | 140.28±466.41 | 2766.61±464.53*** | 3178.48±494.15*** |
| **Total deposit particles/$mm^2$** | 2.91±5.81 | 16.75±5.79 | 39.20±6.03***$^{\#}$ |
| **Total LBs/$mm^2$** | 0.11±1.81 | 3.62±1.80 | 14.47±1.87***$^{\#\#\#}$ |

Univariate tests with covarying age, sex, PMD. Compared to control, *** p<0.001. Compared to stage IV PD, $^{\#}$ p<0.05, $^{\#\#\#}$ p<0.001.

**Supplementary Table 3** Neuronal signal density of PLA and IHC (data presented as mean ± SEM)

| | Controls | PD stage IV | PD stage VI |
|---|---|---|---|
| **PLA particle containing neurons (%)** | 6.76±5.54 | 70.50±5.52*** | 63.51±5.75*** |
| **Deposit particle containing neurons (%)** | 0.50±1.90 | 4.70±1.80 | 10.70±1.90***$^{\#}$ |
| **LB containing neurons (%)** | -0.05±0.59 | 0.87±0.58 | 4.13±0.61***$^{\#\#\#}$ |

Univariate tests with covarying age, sex, PMD. Compared to control, *** p<0.001. Compared to stage IV PD, $^{\#}$ p<0.05, $^{\#\#\#}$ p<0.001.

**Supplementary Table 4** Average PLA particle count in affected neurons (data presented as mean ± SEM)

| | Controls | PD stage IV | PD stage VI |
|---|---|---|---|
| **Superficial grey matter (SGM)** | 1.09±0.58 | 2.09±0.54 | 3.40±0.60* |
| **Deep grey matter (DGM)** | 1.12±0.29 | 2.23±0.24** | 2.18±0.26* |

Univariate tests with covarying age, sex, PMD. Compared to control, * p<0.05, ** p<0.01.

**Supplementary Table 5** Neuronal, extracellular, and glial density of PLA signals in controls versus PD (data presented as mean ± SEM)

| | Category criteria | Controls | PD stage IV | PD stage VI |
|---|---|---|---|---|
| **Proportion of neuronal nuclear type in total neurons (%)** | With ≥ one particle in the neuronal nucleus | 1.67±1.20 | 6.23±1.08** | 5.44±1.11* |
| **Proportion of cytoplasmic type in total neurons (%)** | With ≥ one particle in the neuronal cytoplasm | 6.64±3.98 | 41.08±3.58*** | 30.83±3.68*** |
| **Proportion of mixed type in total neurons (%)** | With ≥ one particle in the neuronal nucleus and cytoplasm | 0.28±4.84 | 23.44±4.36*** | 27.14±4.48*** |
| **Proportion of PLA particle containing neurons (%)** | | 6.76±5.54 | 70.50±5.52*** | 63.51±5.75*** |
| **Extracellular & glial PLA/mm$^2$** | | 63.10±336.93 | 1778.30±335.57*** | 2336.68±349.74*** |
| **Proportion of total PLA particles located in neuronal nuclei & cytoplasm (%)** | | 35.81±8.14 | 33.63±4.62 | 28.92±2.96 |
| **Proportion of total PLA particles located in neuropil/extracellular/glial space (%)** | | 64.19±8.14 | 66.37±4.62 | 71.08±2.96 |

Univariate tests with covarying age, sex, PMD. Compared to control, * p<0.05, ** p<0.01, *** p<0.001.
